# Accelerated single cell seeding in relapsed multiple myeloma

**DOI:** 10.1101/2020.02.25.963272

**Authors:** Heather Landau, Venkata Yellapantula, Benjamin T. Diamond, Even H. Rustad, Kylee H. Maclachlan, Gunes Gundem, Juan Medina-Martinez, Juan Arango Ossa, Max Levine, Yangyu Zhou, Rajya Kappagantula, Priscilla Baez, Marc Attiye, Alvin Makohon-Moore, Lance Zhang, Eileen M Boyle, Cody Ashby, Patrick Blaney, Minal Patel, Yanming Zhang, Ahmet Dogan, David Chung, Sergio Giralt, Oscar B. Lahoud, Jonathan U. Peled, Michael Scordo, Gunjan Shah, Hani Hassoun, Neha S. Korde, Alexander M. Lesokhin, Sydney Lu, Sham Mailankody, Urvi Shah, Eric Smith, Malin L. Hultcrantz, Gary A. Ulaner, Frits van Rhee, Gareth Morgan, C. Ola Landgren, Elli Papaemmanuil, Christine Iacobuzio-Donahue, Francesco Maura

## Abstract

The malignant progression of multiple myeloma is characterized by the seeding of cancer cells in different anatomic sites followed by their clonal expansion. It has been demonstrated that this spatial evolution at varying anatomic sites is characterized by genomic heterogeneity. However, it is unclear whether each anatomic site at relapse reflects the expansion of pre-existing but previously undetected disease or secondary seeding from other sites. Furthermore, genomic evolution over time at spatially distinct sites of disease has not been investigated in a systematic manner.

To address this, we interrogated 25 samples, by whole genome sequencing, collected at autopsy from 4 patients with relapsed multiple myeloma and demonstrated that each site had a unique evolutionary trajectory characterized by distinct single and complex structural variants and copy number changes. By analyzing the landscape of mutational signatures at these sites and for an additional set of 125 published whole exomes collected from 51 patients, we demonstrate the profound mutagenic effect of melphalan and platinum in relapsed multiple myeloma. Chemotherapy-related mutagenic processes are known to introduce hundreds of unique mutations in each surviving cancer cell. These mutations can be detectable by bulk sequencing only in cases of clonal expansion of a single cancer cell bearing the mutational signature linked to chemotherapy exposure thus representing a unique single-cell genomic barcode linked to a discrete time window in each patient’s life. We leveraged this concept to show that multiple myeloma systemic seeding is accelerated at clinical relapse and appears to be driven by the survival and subsequent expansion of a single myeloma cell following treatment with high dose melphalan therapy and autologous stem cell transplant.

## Introduction

The pathogenesis of multiple myeloma is characterized by a long and complex evolutionary process through two clinically defined precursor stages: monoclonal gammopathy of uncertain significance and smoldering multiple myeloma.^1–4^ The progression of precursor disease to invasive multiple myeloma is characterized by branching evolutionary patterns, clonal sweeps together with local evolution and expansion of cancer cells in varying anatomical sites.^5,6^ Divergence and progression at distinct sites of disease - spatial evolution - magnifies the genomic heterogeneity of multiple myeloma, where a range of clones compete for dominance and are positively selected according to their genetic driver landscape reflected in their ability to best adapt to the local environment.^7–12^ Similar evolutionary patterns have also been observed in patients with multiple myeloma at clinical relapse.^13,14^ However, it is unclear if, at the time of relapse, the new disease sites reflect a pre-existing but previously undetected disease localization or a new dissemination of disease “seeding”. Furthermore, despite the unquestionable spatial heterogeneity of multiple myeloma,^13^ to our knowledge, its development over time has not been investigated in a systematic manner.

Somatic mutations in cancer genomes are caused by different mutational processes, each of which generates a characteristic mutational signature.^15^ Considering each single nucleotide variant (SNV) together with its neighboring bases at 5’ and 3’ (the trinucleotide context), more than 40 mutational signatures (or single base signatures; SBS) have been described, some of which are associated with defective DNA repair mechanisms, exposure to exogenous carcinogens, or radical oxygen stress.^16^ Using large genomic datasets, we recently described the landscape of mutational processes active in multiple myeloma.^8,17–21^ At diagnosis (i.e. prior to therapy), the mutational landscape is shaped by seven main mutational processes, five of which have a recognized etiology: activation-induced cytidine deaminase (AID; SBS9), aging (SBS1 and SBS5), and APOBEC (SBS2 and SBS13). At relapse, we reported a new mutational signature associated with melphalan exposure named SBS-MM1.^20^ Similar to other chemotherapy-related mutational signatures described in other malignancies, each surviving myeloma cell exposed to melphalan will acquire a unique set of mutations detectable by bulk sequencing only if a melphalan-exposed single cell is positively selected and expands (**Figure 1A**).^16,22,23^ As a consequence, two different multiple myeloma localizations after melphalan can present three different scenarios. In the first all cancer cells in both localizations will share an identical catalogue of melphalan-related mutations, suggesting that both anatomic sites were seeded by one cancer cell surviving the exposure to melphalan (**Figure 1B**). In the second each localization will have a unique catalogue of melphalan-related mutations, suggesting that cells at the two localizations pre-existed the exposure to melphalan (**Figure 1C**). Finally, it is possible that myeloma cells are reinfused and engraft with the stem cell transplant, avoiding in this way any exposure to melphalan and the associated mutational signature. Although the impact of these chemotherapy-related mutations on cancer aggressiveness at clinical relapse is not fully understood, SBS-MM1 represents unique single-cell genomic barcode for clonal cells derived from a single propagating cell linked to a discrete time point in each patient’s life.

**Figure 1.**
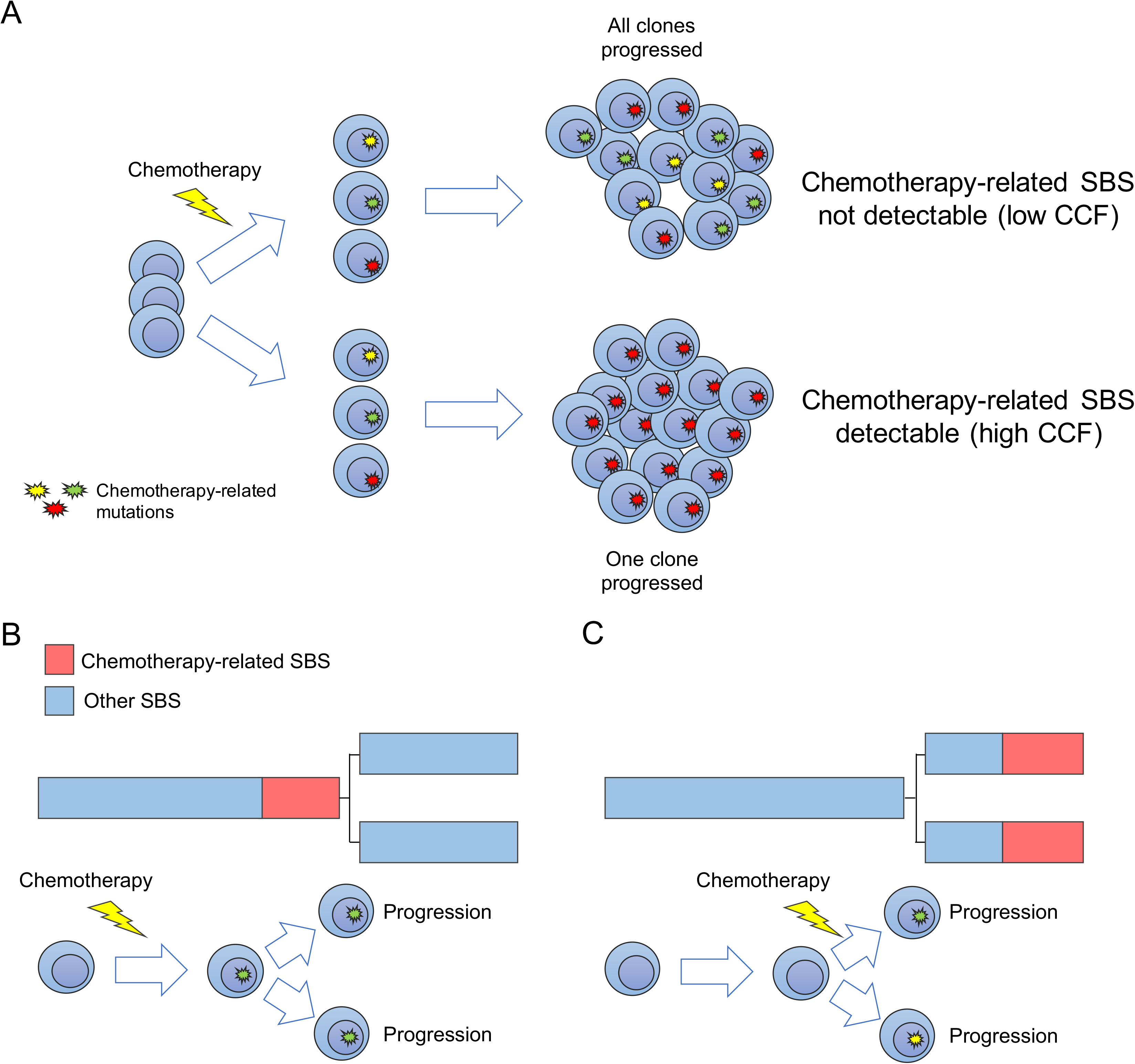
Chemothrapy-related mutational signatures. A) Cartoon summarizing the single cell expansion model. In this model chemotherapy-related mutational signatures will be detectable only if one cancer cell is selected and take the clonal dominance. B-C) Two possible scenario for chemotherapy-related mutational signatures in two different disease localizations (i.e. two different branches of the phylogenetic tree). The phylogenetic tree trunk and branches length represent the (sub)clone mutational load.

Here, to investigate the spatial and temporal systemic dissemination of multiple myeloma at clinical relapse, we interrogated 25 samples collected at warm autopsy from four patients with relapsed/refractory multiple myeloma by whole genome sequencing (WGS). Leveraging the chemotherapy-related mutational signatures caused by melphalan and platinum; we show that disease seeding is accelerated at clinical relapse and appears to be driven by a single myeloma propagating cell that survives after high dose melphalan therapy followed by autologous stem cell transplant.

## Methods

### Patient characteristics

Twenty-one tumor and four non-tumor samples were collected from four patients (**Figure 2A** and **Supplementary Table 1**). All patients consented to autopsy and sample collection as a part of the “Last Wish Program” at Memorial Sloan Kettering Cancer Center (MSKCC).^24^ All patients were treated with multiple lines of therapy (median 6, ranges 5-8) including combinations of novel agents. All patients had previously received at least one round of high dose melphalan therapy followed by autologous stem cell transplant (**Figure 2B**). Two patients had an overall survival longer than 7 years (I-H-106917 and I-H-130719); in contrast, the other two died within 3 years of diagnosis (I-H-130718 and I-H-130720) (**Figure 2B**).

**Figure 2.**
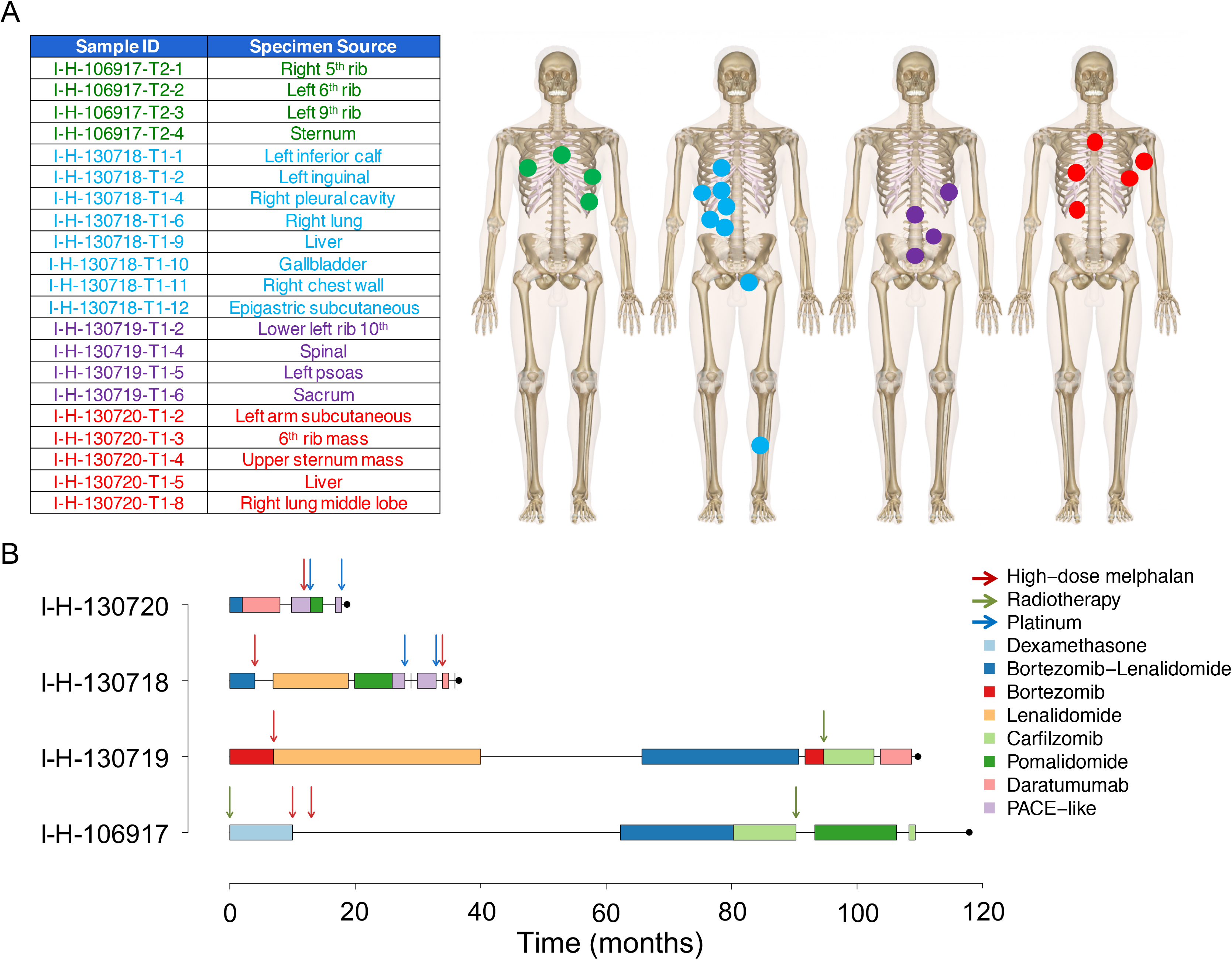
Patient cohort and samples. A) Anatomical sites that were biopsied in each patient. B) Summary of the treatment history of each patient. After the front line, only agents new to each patient were reported. The second I-H-130718 high dose melphalan was the allogeneic stem cell transplant. PACE = cisplatin, doxorubicin, cyclophosphamide and etoposide

All available 18F-fluorodeoxyglucose positron emission tomography/computed tomography (FDG PET/CT) imaging studies for each patient were reviewed by a dual Diagnostic Radiology and Nuclear Medicine board certified radiologist with 15 years of FDG PET/CT experience (G.A.U.) Maximum Intensity Projection images of FDG PET studies and Maximum Standardized Uptake Values (SUVmax) of reference lesions were obtained using PET VCAR (GE Healthcare).

### Sequencing and genomic analysis

Each tumor sample was collected from a different disease localization site (**Figure 2A**) and DNA was extracted from CD138+ purified cells. To avoid contamination related to late systemic disease dissemination, cells collected from skeletal muscles were used as matched normal controls. All normal samples were histologically reviewed to exclude microscopic foci of tumor. Tumor biopsies collected were commonly very cellular and only those with > 70% cellularity based on histologic review were selected for DNA extraction. After PicoGreen quantification and quality control by Agilent BioAnalyzer, 500ng of genomic DNA were sheared using a LE220-plus Focused-ultrasonicator (Covaris catalog # 500569) and sequencing libraries were prepared using the KAPA Hyper Prep Kit (Kapa Biosystems KK8504) with modifications. Briefly, libraries were subjected to a 0.5X size select using aMPure XP beads (Beckman Coulter catalog # A63882) after post-ligation cleanup. Libraries not amplified by PCR (07652_C) were pooled equivolume and were quantitated based on their initial sequencing performance. Libraries amplified with 5 cycles of PCR (07652_D, 07652_F, 07652_G) were pooled equimolar. Samples were run on a NovaSeq 6000 in a 150bp/150bp paired end run, using the NovaSeq 6000 SBS v1 Kit and an S4 flow cell (Illumina).

The median coverage for tumor and normal samples was 92.1X and 58.8X respectively (**Supplementary Table 2**). All bioinformatics analyses were performed using our in-house pipeline Isabl (Medina et al *in preparation*). Briefly, FASTQ files were aligned to the GRCh37 reference genome using BWA-MEM, and de-duplicated aligned BAM files were analyzed using the following published tools: 1) Battenberg for clonal and subclonal copy number aberrations (CNAs);^25^ 2) BRASS for structural variants (SVs);^26^ 3) CaVEMan and Pindel for SNVs and small indels.^27,28^ The clonal composition and phylogenic tree of each multiple myeloma patient was reconstructed by running the Dirichlet Process (DP).^8,25^ Only clusters with more than 50 mutations were considered (as recently described).^8,14^ In the only patient that underwent allogeneic stem cell transplant (I-H-130718), all samples had a subclonal DP cluster of 1594 mutations with a median cancer cell fraction less than 10%, reflecting the unique donor single nucleotide polymorphism (SNP) profile. These mutations were not included in any subsequent analysis.

Complex structural variants such as chromothripsis, templated insertion and chromoplexy were defined as previously reported.^14,29–31^

### Mutational Signatures

Mutational signature analysis was performed applying our recently published workflow, based on three main steps: *de novo* extraction, assignment and fitting.^17^ For the first step, we ran SigProfiler^16^ combining our multi-spatial WGS cohort with a recently published cohort of 52 WGS patients.^9,14,20,32^ Then all extracted signatures were assigned to the latest COSMIC reference (https://cancer.sanger.ac.uk/cosmic/signatures/SBS/) in order to define which known mutational processes were active in our cohort. Finally, we applied our recently developed fitting algorithm (*mmsig*) to confirm the presence and estimate the contribution of each mutational signature in each sample.^20^ Confidence intervals were generated by drawing 1000 mutational profiles from the multinomial distribution, each time repeating the signature fitting procedure, and finally taking the 2.5^th^ and 97.5^th^ percentile for each signature. Mutational signature transcriptional strand bias analysis was performed using SigProfiler and integrated into *mmsig*. The source code of *mmsig* is available on GitHub: https://github.com/evenrus/mmsig.

### Validation set

We imported recently published whole exome sequencing (WXS) data obtained from multiple samples (N=125) collected from different anatomic loci from both newly diagnosed and relapsed multiple myeloma patients (EGAS00001002111, n=51).^13^ In 40 patients, multiple samples were collected at diagnosis from different sites; in the other 11 patients, at least one sample was collected at relapse after intensive treatment, such as platinum-containing regimens and high dose melphalan with autologous stem cell transplant (**Supplementary Table 3-4**). In total, 125 tumor and 51 normal WXS data were included in this study. FASTQ files were aligned to the reference genome using BWA-MEM. BAM files were analyzed for SNV and indels using CaVEMan and Pindel, similarly to the WGS cohort.^27,28^ The CNA profile of each sample was estimated using Facets.^33^ The clonal and subclonal architecture of each patient was reconstructed using the DP for 47 patients.^7^ In 4 patients, the DP failed due to either low CNA quality or the low sample purity. These patients were removed from the study.

## Results

### Phylogenetic tree and disease seeding

To define the key evolutionary trajectories and drivers involved in multiple myeloma systemic seeding, we reconstructed the clonal and subclonal composition of each patient included in the WGS and WXS cohorts (**Methods**). Mutations shared as clonal by all samples from the same individual composed the trunk of the phylogenetic tree. Different late clonal or subclonal clusters could have arisen either directly from the trunk of the phylogenetic tree or from one of its branches. To define the evolutionary history of each patient’s tumor, we reconstructed the most likely phylogenetic tree solution for each patient and defined the main evolutionary trajectories drawing a line from the tip of each branch, via any larger branches and down through the trunk, following the pigeonhole principle (**Figure 3** and **Supplementary Figures 1-2**).^20,34^ A median of 10938 (range 6977-13239) mutations were detected by WGS. The trunk of the two longer-surviving myeloma patients in the WGS cohort accounted for 80.5% and 69% of all mutations. In contrast the trunk was shorter and comprised of far fewer mutations for the two patients with shorter survival (34% and 22% of total mutational burden). This difference may be partially explained by the number of samples from each patient, but also by the major subclonal diversification observed in the patients with a short survival. Interestingly, each disease site in the WGS cohort showed a unique evolutionary trajectory characterized by distinct genomic aberrations (**Figure 3** and **Supplementary Figure 3-4**). The majority of these aberrations were single and complex SVs and CNAs, confirming their critical importance in multiple myeloma progression and evolution.^14,30,35^ In line with previous observations, chromothripsis and templated insertion events were often observed in the trunk, while chromoplexy tended to occur in the branches (**Figure 3**).^14^ In contrast to newly diagnosed multiple myelomas, mutations in known driver genes^19^ were rarely identified in the branches of the phylogenetic tree, and all patients had at least one clone with a mutation involving the RAS pathway.

**Figure 3.**
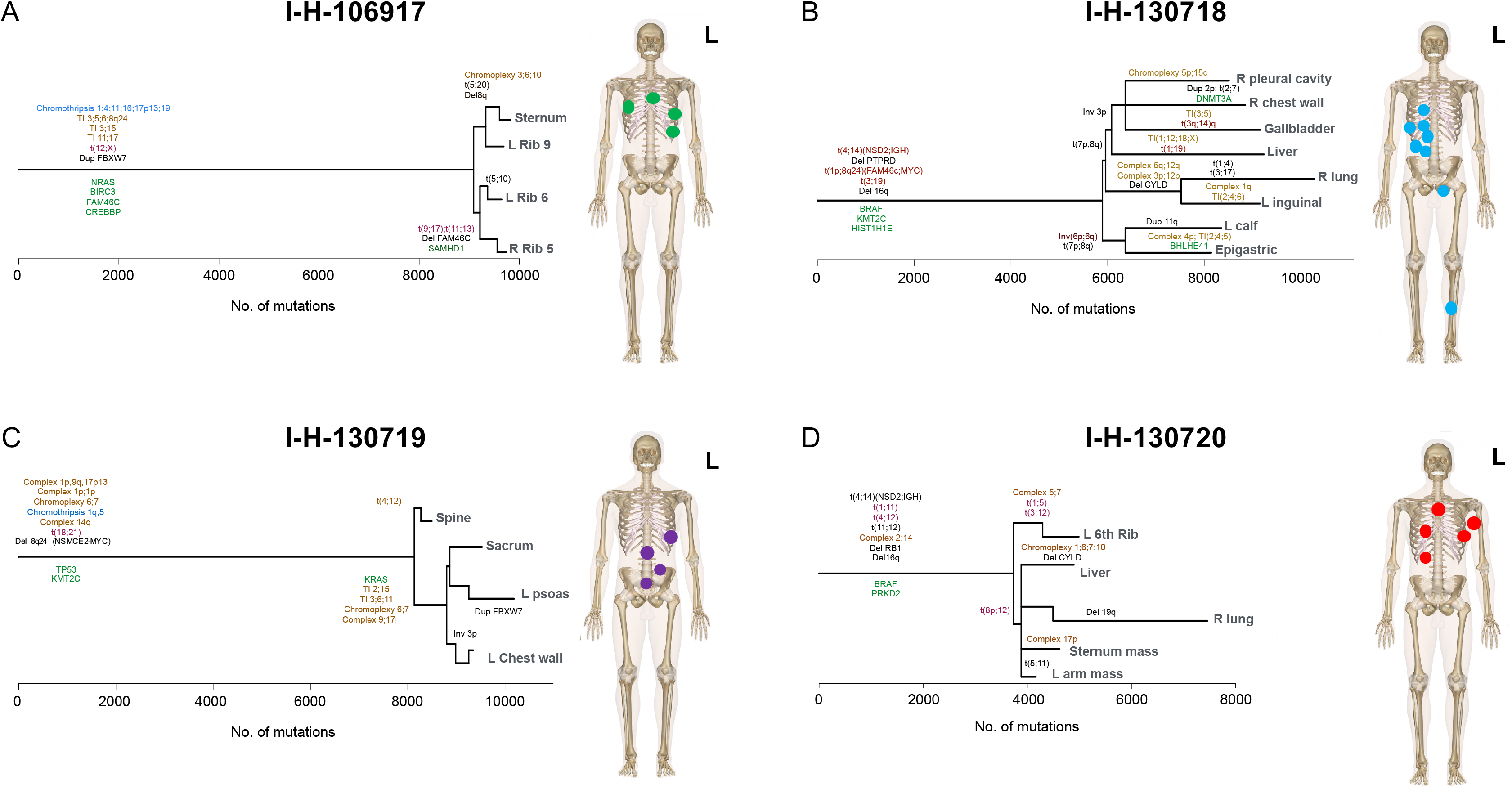
Tumor phylogenies. Phylogenetic trees generated from the Dirichlet process analysis were drawn such that the trunk and branch lengths were proportional to the (sub)clone mutational load. All main drivers (CNAs, SNVs, and SVs) were annotated according to their chronological occurrence and colored according to the type of event. Known driver SNVs were annotated in green, single SVs and CNAs in black, translocations associated with copy number changes in brown, chromothripsis in light blue and other complex events in dark yellow. TI= templated insertion. Lines from different subclone branches are separated by hooks.

Comparing the phylogenetic tree structure and subclonal diversification of the WGS and WXS cohorts, we observed a higher number of evolutionary trajectories in the former. This could be explained by the larger number of specimens from each patient in the WGS series (**Supplementary Figure 5A**), reflecting the high spatial heterogeneity of multiple myeloma detectable in all patients included in this study. Across both series we observed a median of 57 nonsynonymous SNVs per patient (range 21-464). Interestingly, patients with relapsed disease showed a higher number of nonsynonymous SNVs compared with baseline (both WGS and WXS) (**Supplementary Figure 5B**). This higher frequency was independent of the number of subclones and the coverage (which was corrected for sample ploidy and purity) (**Supplementary Figure 5C**).

### Mutational signature landscape

The mutational signature profile of each sample was defined using our recently published workflow.^17^ First, we performed *de novo* extraction of mutational signatures running *SigProfiler* (**Supplementary Figure 6** and **Supplementary Table 5**).^16^ In addition to the 8 known multiple myeloma mutational signatures, we extracted a new signature similar to the recently reported SBS35. This mutational has been shown to be associated with exposure to platinum,^16,22,23,36^ a drug class often included in intensive multiple myeloma chemotherapy regimens. To confirm the presence of each extracted mutational signature and to estimate their contribution to the overall mutational profile, we ran our recently developed fitting algorithm (*mmsig*; **Supplementary Figure 3-4**). In the WGS cohort, SBS-MM1 and its characteristic transcriptional strand bias were observed all patients, consistent with our prior report.^30^ As expected SBS35 was identified only in the two patients who received a platinum-based treatment (I-H-130718 and I-H-130720). In the WXS cohort, SBS35 and SBSMM1 were observed only among mutations acquired or selected after treatment (**Supplementary Figure 7**). These data suggest that, similarly to other cancers, the mutational landscape of clinically relapsed multiple myeloma is heavily shaped by exposure to distinct chemotherapeutic agents, such as melphalan or platinum.

To investigate the impact of these chemotherapy-related mutations on the multiple myeloma genomic profile, we combined the WGS and WXS cohorts and estimated the contribution of both SBS35 and SBS-MM1 among nonsynonymous SNVs.^23^ Interestingly, 25.7% (CI 95% 20-32%) of all nonsynonymous mutations at clinical relapse were caused by one of these two mutational processes suggesting that chemotherapy exposure might play a role in increasing genomic complexity that may associate clinically aggressive disease at relapse (**Figure 4**).

**Figure 4.**
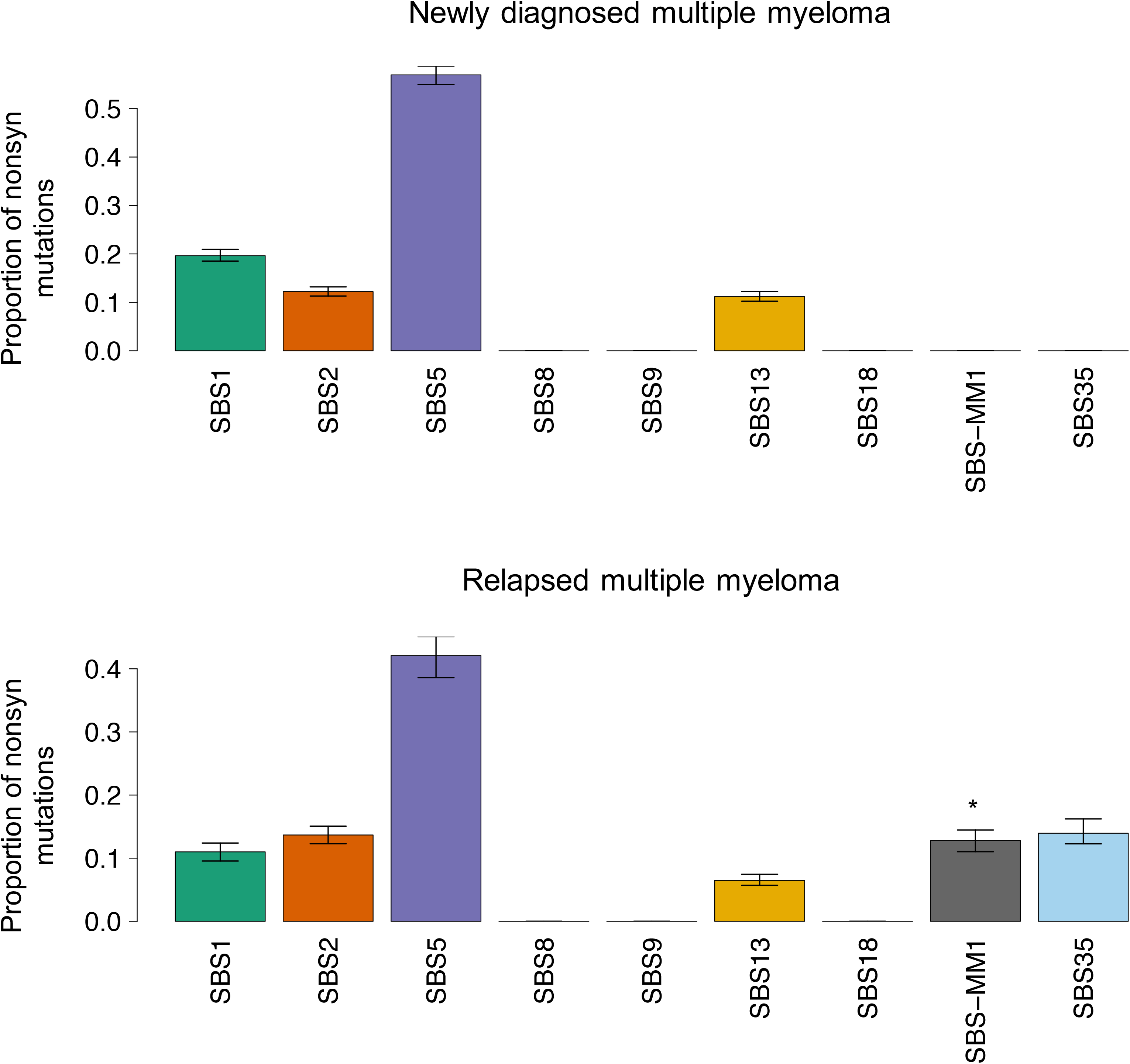
The mutational signature landscape of nonsynonymous (nonsyn) mutations at diagnosis (A) and relapse (B). The asterisk reflects the presence of transcriptional strand bias for SBS-MM1.

To reconstruct the timeline of mutational processes for each patient, we ran *mmsig* on each DP cluster of mutations (**Figure 5A**). In line with its activity early in disease development, SBS9 (AID) was detected mostly in the trunk of the phylogenetic trees, while APOBEC activity was detected in both clonal and subclonal clusters.^4,8,19,20^ SBS35 was only detected in the branches of the two post-platinum relapsed patients from WGS. SBS-MM1 showed a heterogenous landscape. In I-H-106917, SBS-MM1 was detected in the trunk and in all first level branches, but not in the second and third level ones (**Figure 5B**). This profile is consistent with a melphalan signature common to all cells, with another subset of melphalan-induced mutations accrued from a second exposure. In I-H-130719, all SBS-MM1 mutations were assigned to the trunk, consistent with exposure to high dose melphalan therapy followed by autologous stem cell transplant received as part of the administered front-line therapy. I-H-130718 was one of the two cases with short survival and platinum exposure. SBS-MM1 was detected in the trunk and in all the latest branches, reflecting the front-line high dose melphalan therapy followed by autologous stem cell transplant as well as the allogeneic stem-cell transplant with melphalan-containing conditioning being used at clinical relapse (**Figure 5C**). In these 3 patients, SBS-MM1 was detected in the trunk, consistent with the expansion and the clonal dominance of a single cell surviving the melphalan-exposure. In fact, the large number of SBS-MM1 related mutations shared by all different sites can only be explained by the existence of a recent common ancestor selected after high dose melphalan therapy followed by autologous stem cell transplant (**Figure 1A-B**). These data also show that in these 3 patients, differences between anatomic sites were not associated with pre-existing undetectable subclones, but by the dissemination from a single multiple myeloma propagating cell that had survived high dose melphalan therapy followed by autologous stem cell transplant. This model is also supported by available PET/CT imaging data, that demonstrates during serial relapses that the majority of end-stage lesions were not detectable until the last progression event before death and subsequent postmortem examination (**Figure 6**; **Supplementary Table 6**).

**Figure 5.**
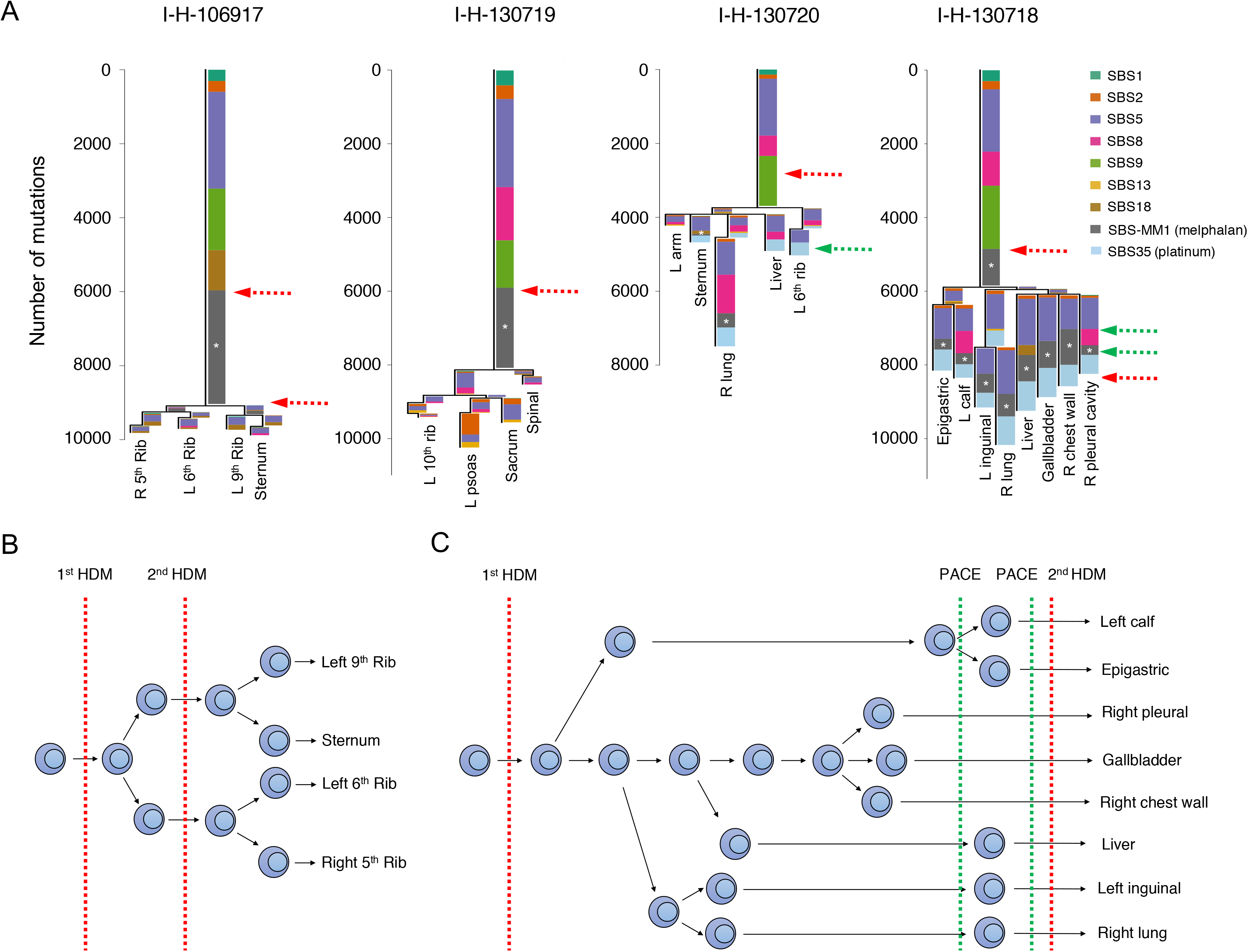
Timeline of mutational signatures. A) Mutational signatures’ contribution for each phylogenetic tree cluster. Asterisks indicates the presence of transcriptional strand bias for SBS-MM1. Green and red dashed arrows represent exposure to platinum and melphalan therapies, respectively. B-C) Cartoon summarizing the relationship between chemotherapy, subclonal selection and seeding in I-H-106917 and I-H-130718. HDM = high dose melphalan, PACE = cisplatin, doxorubicin, cyclophosphamide and etoposide. Green and red dashed lines represent exposure to platinum and melphalan therapies, respectively.

**Figure 6.**
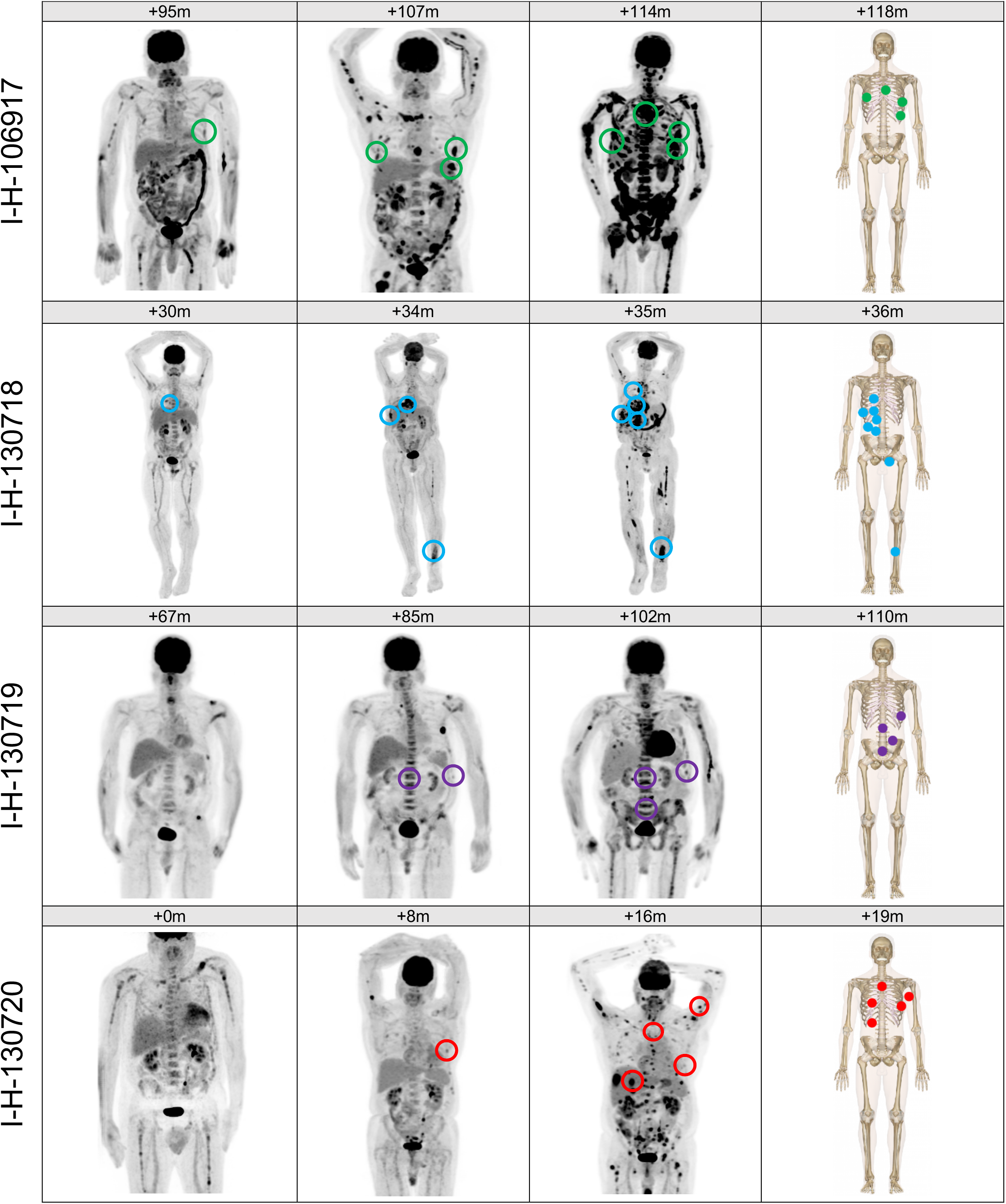
Tracking of biopsied lesions through the disease course by FDG PET. Biopsy sites and FDG PET correlates on Maximum Intensity Projection images for each patient arranged by the number of months following diagnosis (m=months). All included scans show phases of active disease. Biopsy sites were only annotated on FDG PET if identified on radiology review.

In I-H-130718, the time lag between the first and the second round of high dose melphalan therapy was 25 months (**Figure 2B**). In this short time window, we show that a single MM propagating cell survived exposure to high dose melphalan and its subsequent dissemination drove relapse (**Figure 5** and **6**). Then, the exposure of cells at these diverse sites when treated with a platinum-containing regimen and high dose melphalan, increased their mutational burden (**Figure 5C** and **Supplementary Figure 4**).

The systemic seeding in I-H-130720 did not fit in the above described single-cell expansion model, having detectable SBS-MM1 only in two out of five branches. This distribution together with the short survival and relapsed/refractory disease might reflect either the absence of a single cell expansion post-melphalan in some pre-existing disease localizations or the engraftment of clones re-infused with the autologous stem cell transplant.^20^

Overall, these data suggest that a single cell has the potential to drive disease progression, and that systemic seeding of a single clonal cell can occur rapidly at relapse. To further investigate this hypothesis and to explore potential differences between pre- and post-treatment disease seeding, we quantified the differential contribution of mutational signatures associated with aging (SBS1 and SBS5, described as “clock-like”) between the branches and the trunk.^37,38^ The SBS5 profile has significant overlap with both SBS-MM1 and SBS35, and this can lead to the incorrect assignment of signatures, one of the major issues in mutational signature analysis.^17^ To avoid this, we focused on the ratio of SBS1 between branches and the trunk in each patient included in the WGS and the WXS cohorts. The ratio for each patient was corrected for the number of evolutionary trajectories, avoiding the pooling of mutations that were acquired in parallel. Interestingly, the SBS1 ratios were significantly higher in the treatment-naïve patients compared to those observed in the relapsed WXS and WGS cases (**Figure 7A** and **Supplementary Figure 8**), consistent with a subclonal diversification having occurred over a long period of time. These findings support the model in which multiple myeloma seeding can be accelerated following high dose treatment in comparison to that seen during spontaneous evolution.

**Figure 7.**
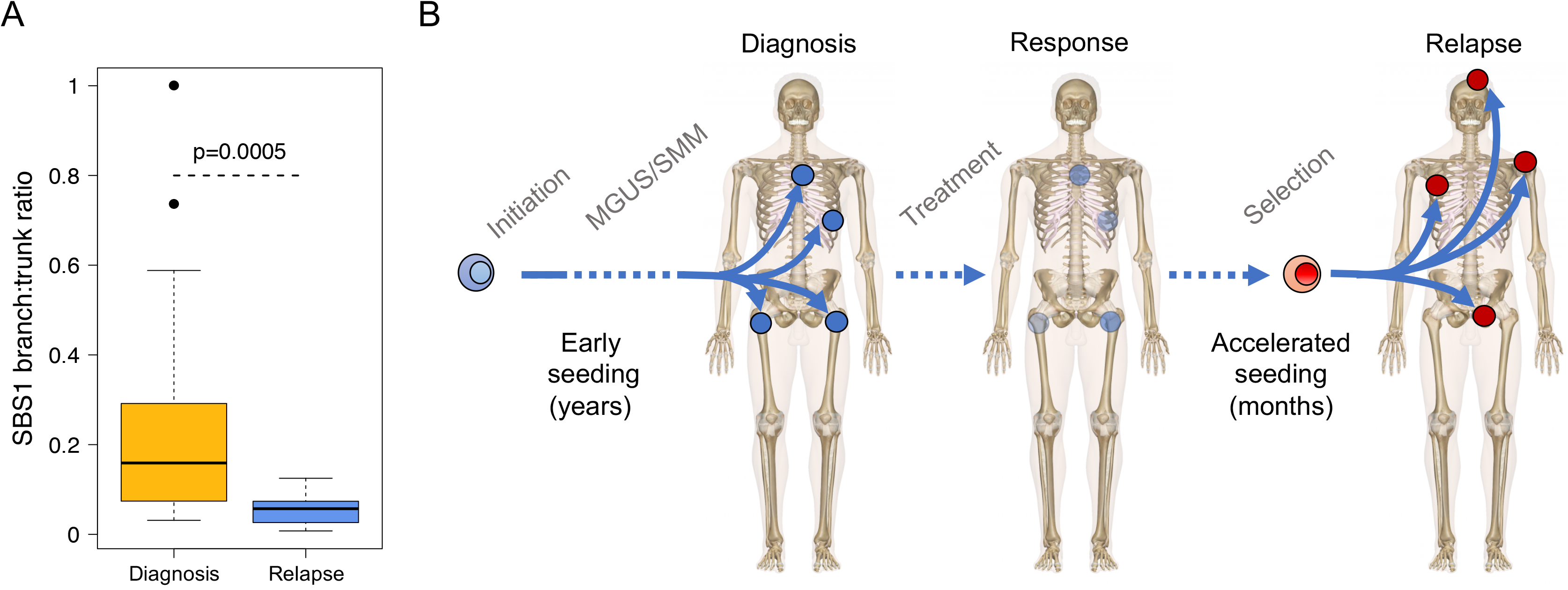
Early versus late divergence of disease sites. A) SBS1 is a mutational signature reflective of biological cell aging. By estimating changes in SBS1 in the trunk and the branches, we were able to study the speed of the tumor evolution in a given site at a given time-point. Here, we show the difference in SBS1 branch:trunk ratio between newly diagnosed and relapsed multiple myeloma (p-value estimated using Wilcoxon test). A Lower SBS1 branch:trunk ratio represents a highly accelerated evolution since divergence from the trunk (most recent common ancestor). B) Cartoon summarizing the seeding patterns over time, starting as slow seeding and tumor growth during the precursor phase, with acceleration in advanced disease

## Discussion

In this study, we used multiple concurrent samples obtained from several rarely biopsied anatomical sites, obtained by warm autopsy in patients with relapsed/refractory multiple myeloma. This unique sample source and study design allowed us to interrogate the spatio-temporal genomic heterogeneity of multiple myeloma at the time of aggressive relapse in a set of patients previously exposed to high dose melphalan. Investigating the mutational signature landscape, we showed the strong mutagenic activity of melphalan and platinum on the multiple myeloma propagating cell and its contribution to the post relapse genomic landscape. While these mutations tend to occur in the late-replicating and non-coding parts of the genome,^20^ we provide evidence that exposure to chemotherapy is responsible for a considerable proportion (>20%) of the nonsynonymous mutations acquired at clinical relapse, with potential effects on subsequent evolutionary trajectories at the time of aggressive clinical relapse.

We demonstrate that multiple myeloma seeding is promoted by an evolutionary process in which distinct clones harboring distinct drivers are selected and expanded at varying anatomic sites. Using chemotherapy-related mutational signatures (i.e. melphalan and platinum) as a genomic barcode, we demonstrate that how this complex process can be driven by a single surviving cell, potentially able to disseminate throughout the entire body. Linking these mutational signatures with the documented timing of chemotherapy exposure, we showed that, at clinical relapse, systemic seeding of multiple myeloma can occur in a very short time window. Importantly, the patterns we demonstrate at relapse are strikingly different to the spatio-temporal patterns of evolution that have been demonstrated during spontaneous evolution prior to the time of initial diagnosis and exposure to therapy. In fact, at diagnosis, different anatomic disease sites are characterized by high burden of clock-like mutation (i.e. SBS1), consistent with an early divergence from the most common recent ancestor followed by slow growth (**Figure 7B**). The accelerated anatomic dissemination we describe at relapse is similar to the metastatic seeding recently reported in different solid cancers.^38,39^ This process is likely the result of a combination of two factors: (1) the selection of a more aggressive/proliferative clones, and, (2) treatment-related immunosuppression. While the first factor is often unpredictable and undetectable with the current bulk sequencing technologies, the second factor represents a rational approach to improve treatment efficacy. Indeed, strategies encouraging immune reconstitution may prevent this acceleration and reduce the incidence of clinical relapse.

These data highlight the importance of considering the complex spatial and temporal heterogeneity of multiple myeloma in the evaluation of treatment-response and in minimal residual disease assessment and provide a strong rationale for comprehensive characterization of the multiple myeloma genomic complexity to enhance clinical decision making.

## Acknowledgements

This work is supported by the Memorial Sloan Kettering Cancer Center NCI Core Grant (P30 CA 008748).

F.M. is supported by the American Society of Hematology, the International Myeloma Foundation and The Society of Memorial Sloan Kettering Cancer Center.

K.H.M. is supported by the Haematology Society of Australia and New Zealand New Investigator Scholarship and the Royal College of Pathologists of Australasia Mike and Carole Ralston Travelling Fellowship Award.

G.J.M is supported by The Leukemia Lymphoma Society.

C.I.D. is supported by NCI grants R35 CA220508 and U2C CA233284 and the Kleberg Foundation.

J.U.P reports funding from NHLBI NIH Award K08HL143189 and the Parker Institute for Cancer Immunotherapy at Memorial Sloan Kettering Cancer Center.

## Author contributions

HL, FM, EP, CID, OL designed and supervised the study, collected and analyzed data and wrote the paper. VY collected and analyzed data and wrote the paper. BTD, HER, KHM, GG, JMM, JAO, ML, JZ analyzed data.

RK, PB, MA, AMM, LZ, MP, EB, CA, PB, YZ, AD, DC, SG, OBL, JUP, MS, GS, HH, MLH, AML, SL, NSK, SM, ES, US, PB, MA, LZ, FVR, GAU and GM collected the data.

## References

1. Corre J, Munshi N, Avet-Loiseau H. Genetics of multiple myeloma: another heterogeneity level? Blood. 2015;125(12):1870–1876.

2. Manier S, Salem KZ, Park J, Landau DA, Getz G, Ghobrial IM. Genomic complexity of multiple myeloma and its clinical implications. Nat Rev Clin Oncol. 2017;14(2):100–113.

3. Morgan GJ, Walker BA, Davies FE. The genetic architecture of multiple myeloma. Nat Rev Cancer. 2012;12(5):335–348.

4. Maura F, Bolli N, Rustad EH, Hultcrantz M, Munshi N, Landgren O. Moving From Cancer Burden to Cancer Genomics for Smoldering Myeloma: A Review. JAMA Oncol. 2019.

5. Rajkumar SV, Landgren O, Mateos MV. Smoldering multiple myeloma. Blood. 2015;125(20):3069–3075.

6. Rajkumar SV, Dimopoulos MA, Palumbo A, et al. International Myeloma Working Group updated criteria for the diagnosis of multiple myeloma. Lancet Oncol. 2014;15(12):e538–548.

7. Bolli N, Avet-Loiseau H, Wedge DC, et al. Heterogeneity of genomic evolution and mutational profiles in multiple myeloma. Nat Commun. 2014;5:2997.

8. Bolli N, Maura F, Minvielle S, et al. Genomic patterns of progression in smoldering multiple myeloma. Nat Commun. 2018;9(1):3363.

9. Lohr JG, Stojanov P, Carter SL, et al. Widespread genetic heterogeneity in multiple myeloma: implications for targeted therapy. Cancer Cell. 2014;25(1):91–101.

10. Walker BA, Wardell CP, Melchor L, et al. Intraclonal heterogeneity is a critical early event in the development of myeloma and precedes the development of clinical symptoms. Leukemia. 2014;28(2):384–390.

11. Bustoros M, Park J, Salem K, et al. Next Generation Sequencing Identifies Smoldering Multiple Myeloma Patients with a High Risk of Disease Progression. Blood. 2017;130 (Supplement 1): 392.

12. Misund K, Keane N, Stein CK, et al. MYC dysregulation in the progression of multiple myeloma. Leukemia. 2020;34(1):322–326.

13. Rasche L, Chavan SS, Stephens OW, et al. Spatial genomic heterogeneity in multiple myeloma revealed by multi-region sequencing. Nat Commun. 2017;8(1):268.

14. Maura F, Bolli N, Angelopoulos N, et al. Genomic landscape and chronological reconstruction of driver events in multiple myeloma. Nat Commun. 2019;10(1):3835.

15. Alexandrov LB, Nik-Zainal S, Wedge DC, et al. Signatures of mutational processes in human cancer. Nature. 2013;500(7463):415–421.

16. Alexandrov LB, Kim J, Haradhvala NJ, et al. The repertoire of mutational signatures in human cancer. Nature. 2020;578(7793):94–101.

17. Maura F, Degasperi A, Nadeu F, et al. A practical guide for mutational signature analysis in hematological malignancies. Nat Commun. 2019;10(1):2969.

18. Maura F, Petljak M, Lionetti M, et al. Biological and prognostic impact of APOBEC-induced mutations in the spectrum of plasma cell dyscrasias and multiple myeloma cell lines. Leukemia. 2017.

19. Maura F, Rustad EH, Yellapantula V, et al. Role of AID in the temporal pattern of acquisition of driver mutations in multiple myeloma. Leukemia. 2019.

20. Rustad H, Yellapantula V, Bolli N, et al. Timing the Initiation of Multiple Myeloma. Sneak Peek. 2019.

21. Walker BA, Wardell CP, Murison A, et al. APOBEC family mutational signatures are associated with poor prognosis translocations in multiple myeloma. Nat Commun. 2015;6:6997.

22. Kucab JE, Zou X, Morganella S, et al. A Compendium of Mutational Signatures of Environmental Agents. Cell. 2019;177(4):821–836 e816.

23. Pich O, Muinos F, Lolkema MP, Steeghs N, Gonzalez-Perez A, Lopez-Bigas N. The mutational footprints of cancer therapies. Nat Genet. 2019;51(12):1732–1740.

24. Iacobuzio-Donahue CA, Michael C, Baez P, Kappagantula R, Hooper JE, Hollman TJ. Cancer biology as revealed by the research autopsy. Nat Rev Cancer. 2019;19(12):686–697.

25. Nik-Zainal S, Van Loo P, Wedge DC, et al. The life history of 21 breast cancers. Cell. 2012;149(5):994–1007.

26. Nik-Zainal S, Davies H, Staaf J, et al. Landscape of somatic mutations in 560 breast cancer whole-genome sequences. Nature. 2016;534(7605):47–54.

27. Jones D, Raine KM, Davies H, et al. cgpCaVEManWrapper: Simple Execution of CaVEMan in Order to Detect Somatic Single Nucleotide Variants in NGS Data. Curr Protoc Bioinformatics. 2016;56:15 10 11–15 10 18.

28. Raine KM, Hinton J, Butler AP, et al. cgpPindel: Identifying Somatically Acquired Insertion and Deletion Events from Paired End Sequencing. Curr Protoc Bioinformatics. 2015;52:15 17 11–12.

29. Korbel JO, Campbell PJ. Criteria for inference of chromothripsis in cancer genomes. Cell. 2013;152(6):1226–1236.

30. Rustad E, Yellapantula V, Glodzik D, et al. Revealing the impact of recurrent and rare structural variants in multiple myeloma. Bioerxiv. 2019.

31. Li Y, Roberts ND, Wala JA, et al. Patterns of somatic structural variation in human cancer genomes. Nature. 2020;578(7793):112–121.

32. Chapman MA, Lawrence MS, Keats JJ, et al. Initial genome sequencing and analysis of multiple myeloma. Nature. 2011;471(7339):467–472.

33. Shen R, Seshan VE. FACETS: allele-specific copy number and clonal heterogeneity analysis tool for high-throughput DNA sequencing. Nucleic Acids Res. 2016;44(16):e131.

34. Mitchell TJ, Turajlic S, Rowan A, et al. Timing the Landmark Events in the Evolution of Clear Cell Renal Cell Cancer: TRACERx Renal. Cell. 2018;173(3):611–623 e617.

35. Barwick BG, Neri P, Bahlis NJ, et al. Multiple myeloma immunoglobulin lambda translocations portend poor prognosis. Nat Commun. 2019;10(1):1911.

36. Boot A, Huang MN, Ng AWT, et al. In-depth characterization of the cisplatin mutational signature in human cell lines and in esophageal and liver tumors. Genome Res. 2018;28(5):654–665.

37. Alexandrov LB, Jones PH, Wedge DC, et al. Clock-like mutational processes in human somatic cells. Nat Genet. 2015;47(12):1402–1407.

38. Noorani A, Li X, Goddard M, et al. Genomic evidence supports a clonal diaspora model for metastases of esophageal adenocarcinoma. Nat Genet. 2020;52(1):74–83.

39. Rabbie R, Ansari-Pour N, Cast O, et al. Multi-site clonality analyses uncovers pervasive subclonal heterogeneity and branching evolution across melanoma metastases. bioarxiv. 2019.

