## Supplementary Material for "Accelerated single cell seeding in relapsed multiple myeloma"

### Supplementary figures

**Supplementary Figure 1.** Phylogenetic tree reconstruction of each newly diagnosed patient included in the WXS cohort.

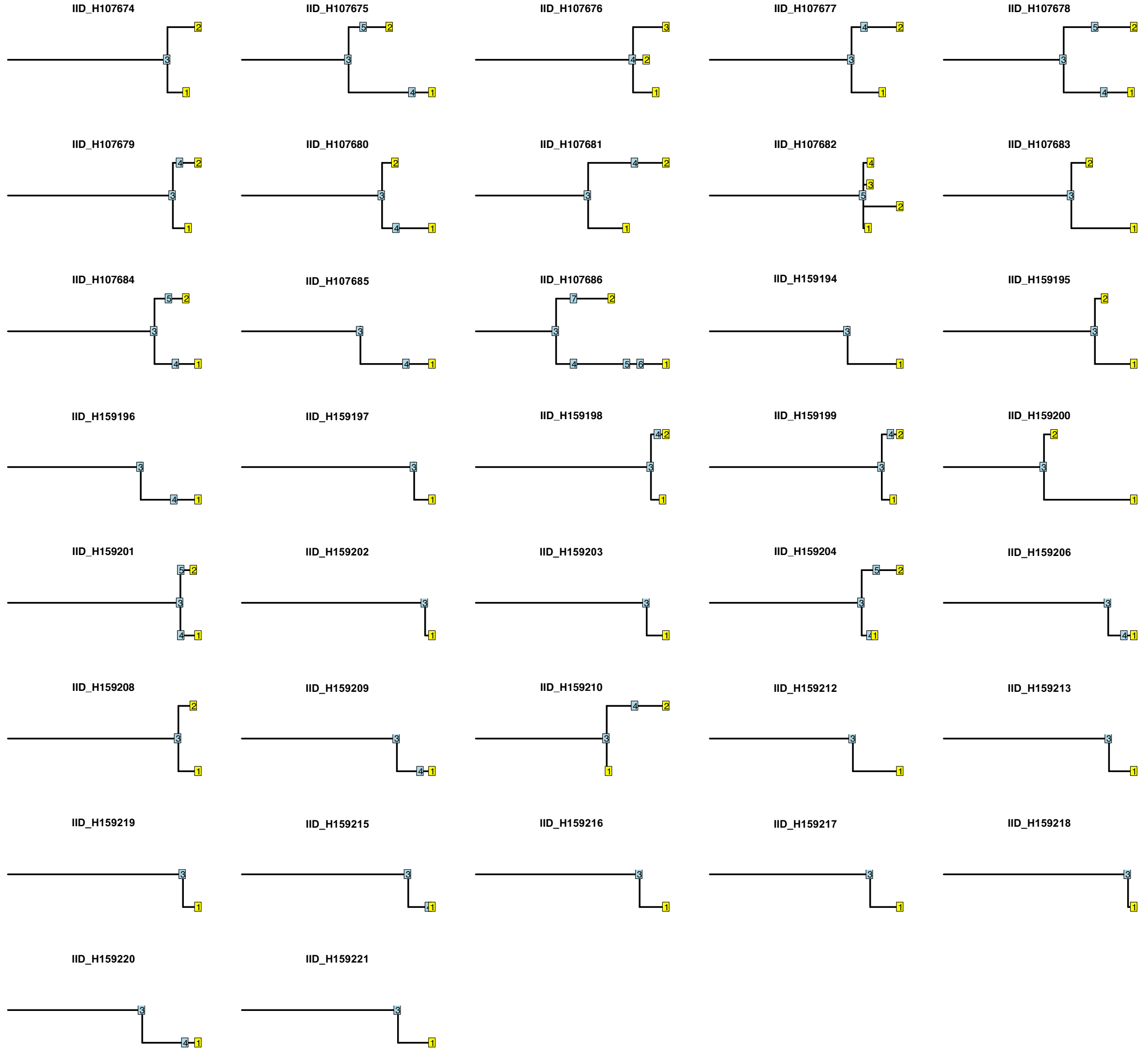

**Supplementary Figure 2.** Phylogenetic tree reconstruction of all patients with at least one sample collected at relapse included in the WXS cohort.

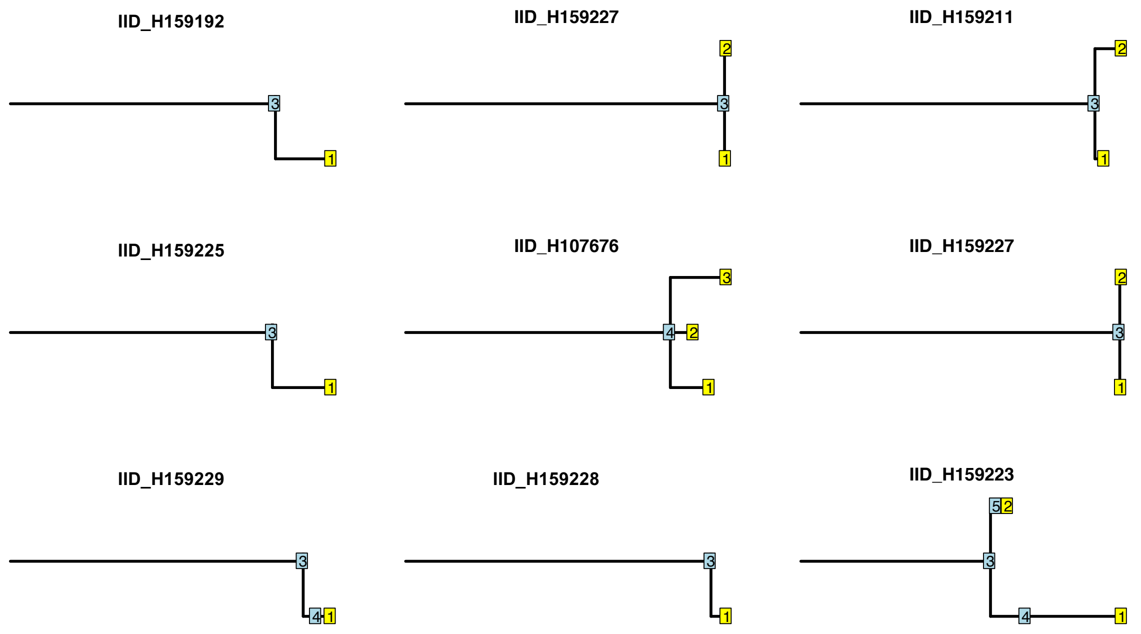

**Supplementary Figure 3.** Genome plot and mutational signatures' landscape of the two patients in the WGS cohort with long survival. The plot on the right showed all the genomic events shared by all samples (events in the trunk of the phylogenetic tree). The plot on the left showed the events not shared by all samples (i.e. events in the branches of the phylogenetic tree). Copy number aberrations are annotated in the circus plot (blue= gain; red = loss of heterozygosity). Structural variants are reported within the circle (black = translocations; blue = inversions; green = tandem-duplications; red= deletions). The asterisk in the mutational signatures barplot reflects the presence of transcriptional strand bias for SBS-MM1.

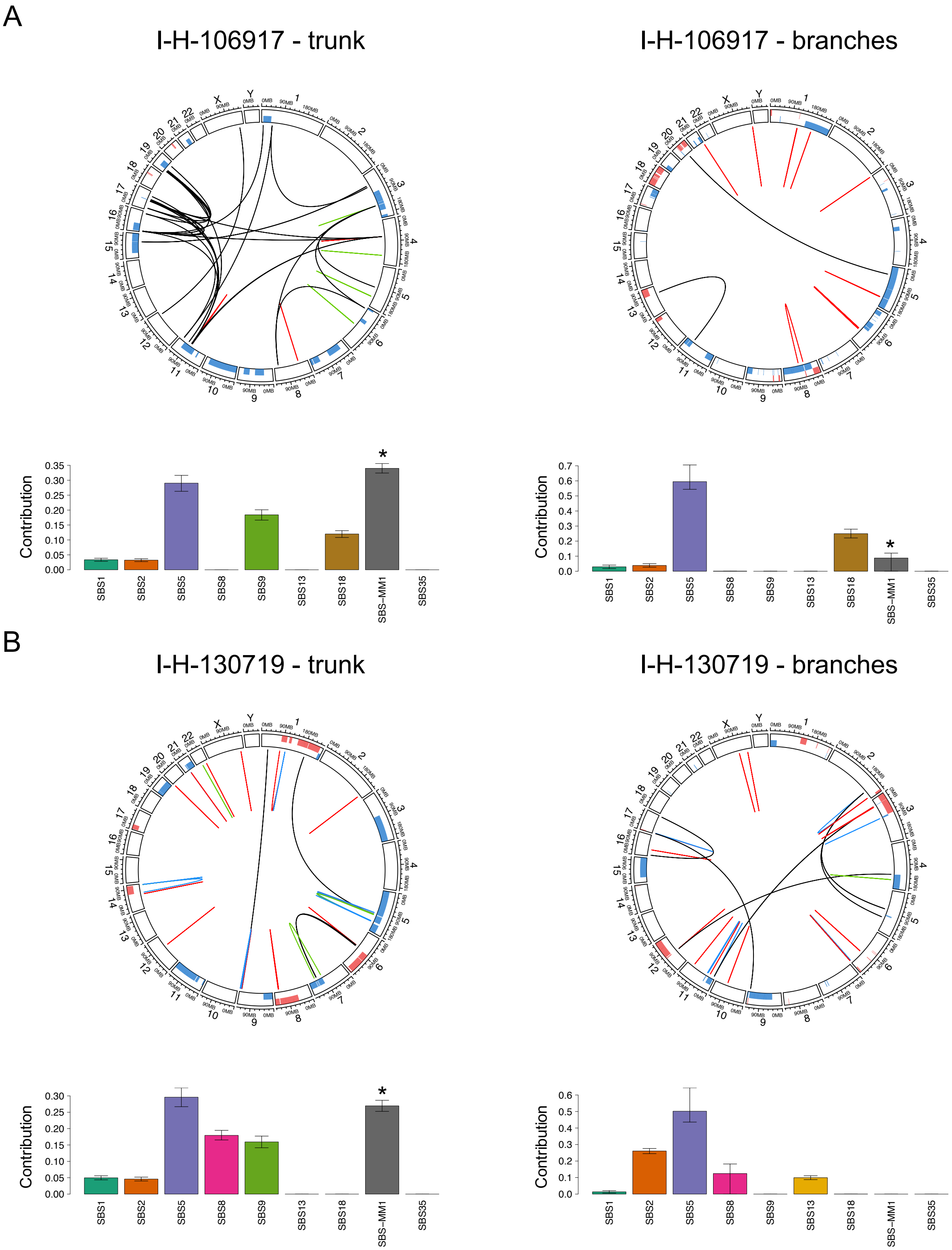
A
I-H-106917 - trunk
I-H-106917 - branches
Contribution
SBS1
SBS2
SBS5
SBS8
SBS9
SBS13
SBS18
SBS−MM1
SBS35
B
I-H-130719 - trunk
I-H-130719 - branches

**Supplementary Figure 4.** Genome plot of the two patients in the WGS cohort with short survival. The plot on the right showed all the genomic events shared by all samples (events in the trunk of the phylogenetic tree). The plot on the left showed the events not shared by all samples (i.e. events in the branches of the phylogenetic tree). Copy number aberrations are annotated in the circus plot (blue= gain; red = loss of heterozygosity). Structural variants are reported within the circle (black = translocations; blue = inversions; green = tandem-duplications; red= deletions). The asterisk in the mutational signatures barplot reflects the presence of transcriptional strand bias for SBS-MM1.

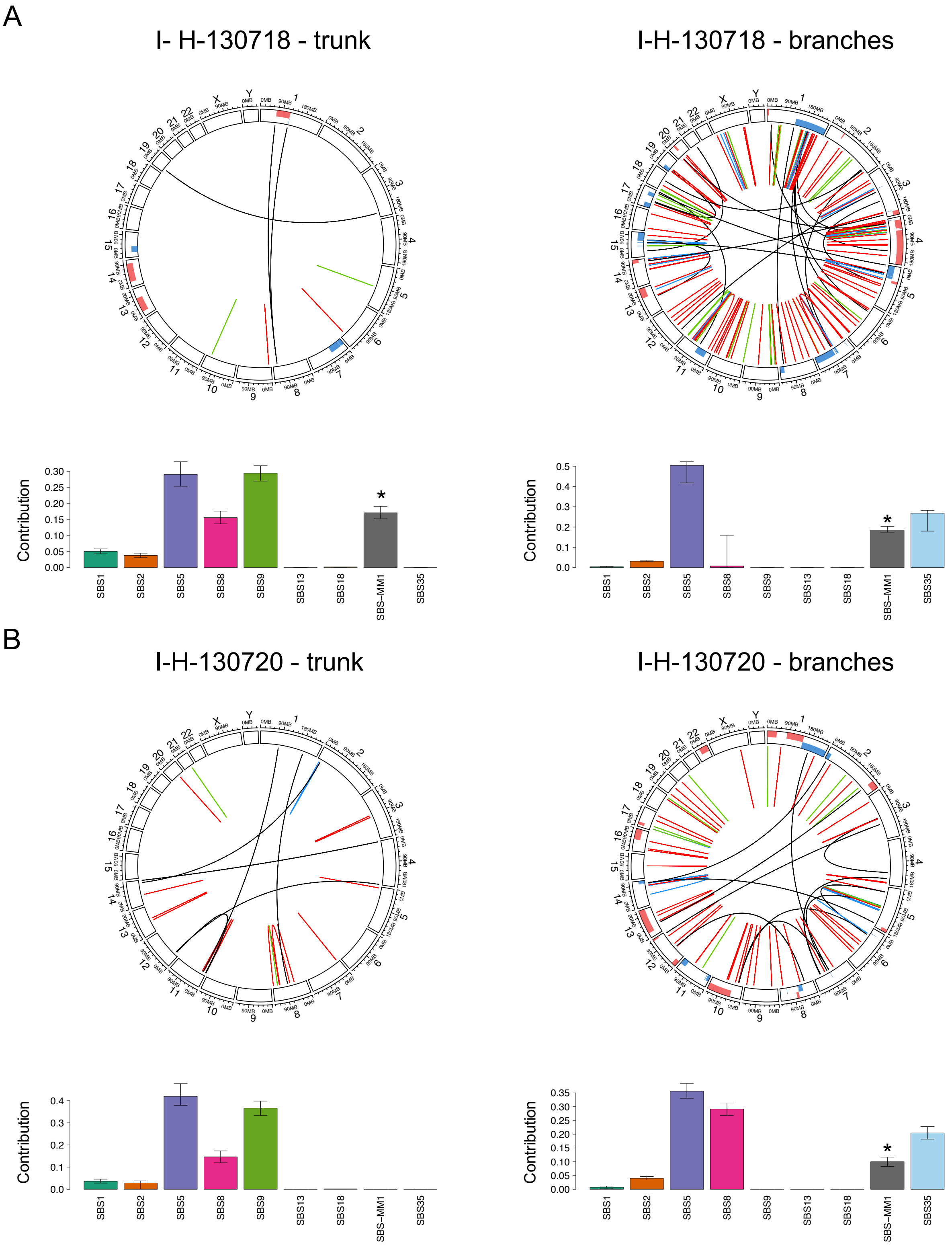
A
I- H-130718 - trunk
I-H-130718 - branches
Contribution
SBS1
SBS2
SBS5
SBS8
SBS9
SBS13
SBS18
SBS−MM1
SBS35
B
I-H-130720 - trunk
I-H-130720 - branches

**Supplementary Figure 5.** Increased burden of nonsynonymous SNVs in relapsed multiple myeloma. A) Linear regression between the number of samples and evolutionary trajectories (number of clones). Blue, red and yellow dots represent relapsed WGS, relapsed WXS and newly diagnosed WXS samples, respectively. B) Boxplot showing increased burden of nonsynonymous SNVs in relapsed multiple myeloma. p-values were estimated using Wilcoxon test. WGS RR = whole genome sequencing at relapsed, WXS DG = whole exome sequencing at diagnosis, WXS RR = whole exome sequencing at relapsed. C) Relative contribution to the R square in a multivariate linear regression model shown in Figure 3A. Estimates were generated using the *relaimpo* R package.

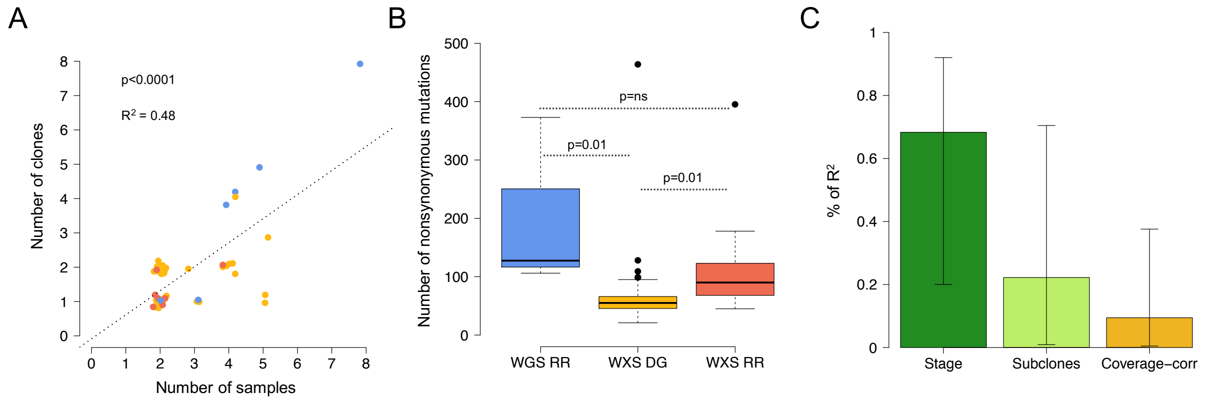

**Supplementary Figure 6.** The 8 mutational signatures extracted by SigProfiler. On the top left of each 96-mutational profile is annotated the signature(s) that contribute to the profile.

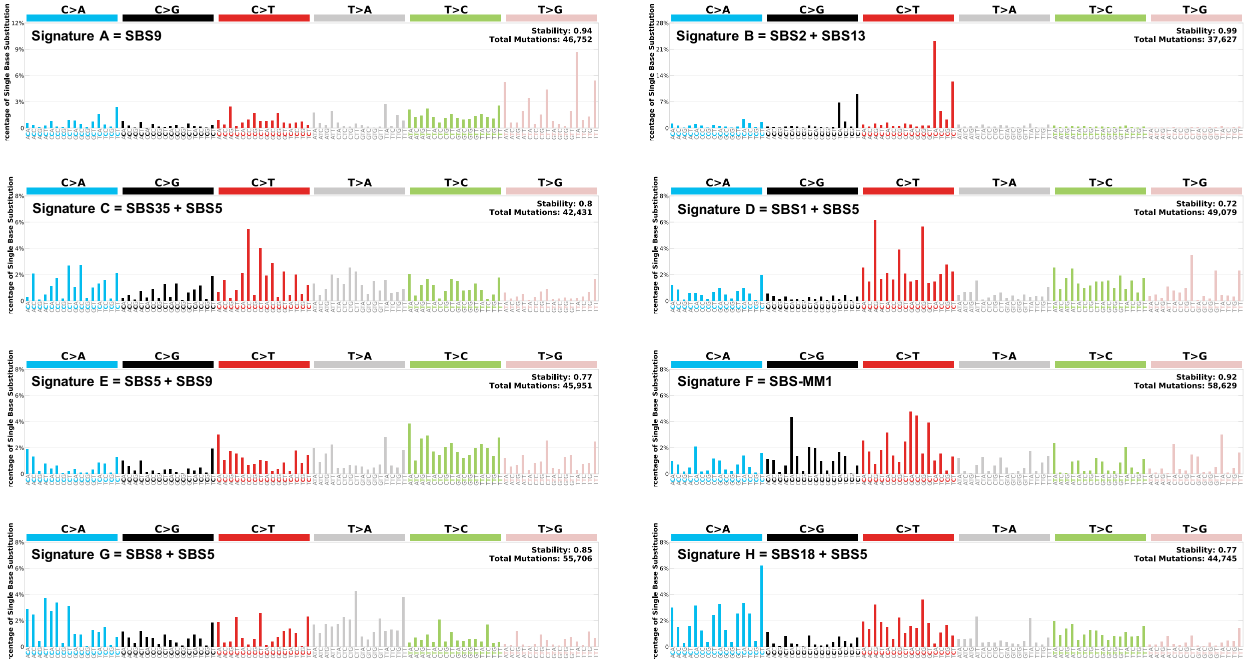

**Supplementary Figure 7.** The 96-mutational profile and mutational contribution of the trunks and the branches of all newly diagnosed and relapsed multiple myeloma included in the WXS cohort. The asterisk in d reflect the presence of transcriptional strand bias for SBS-MM1.

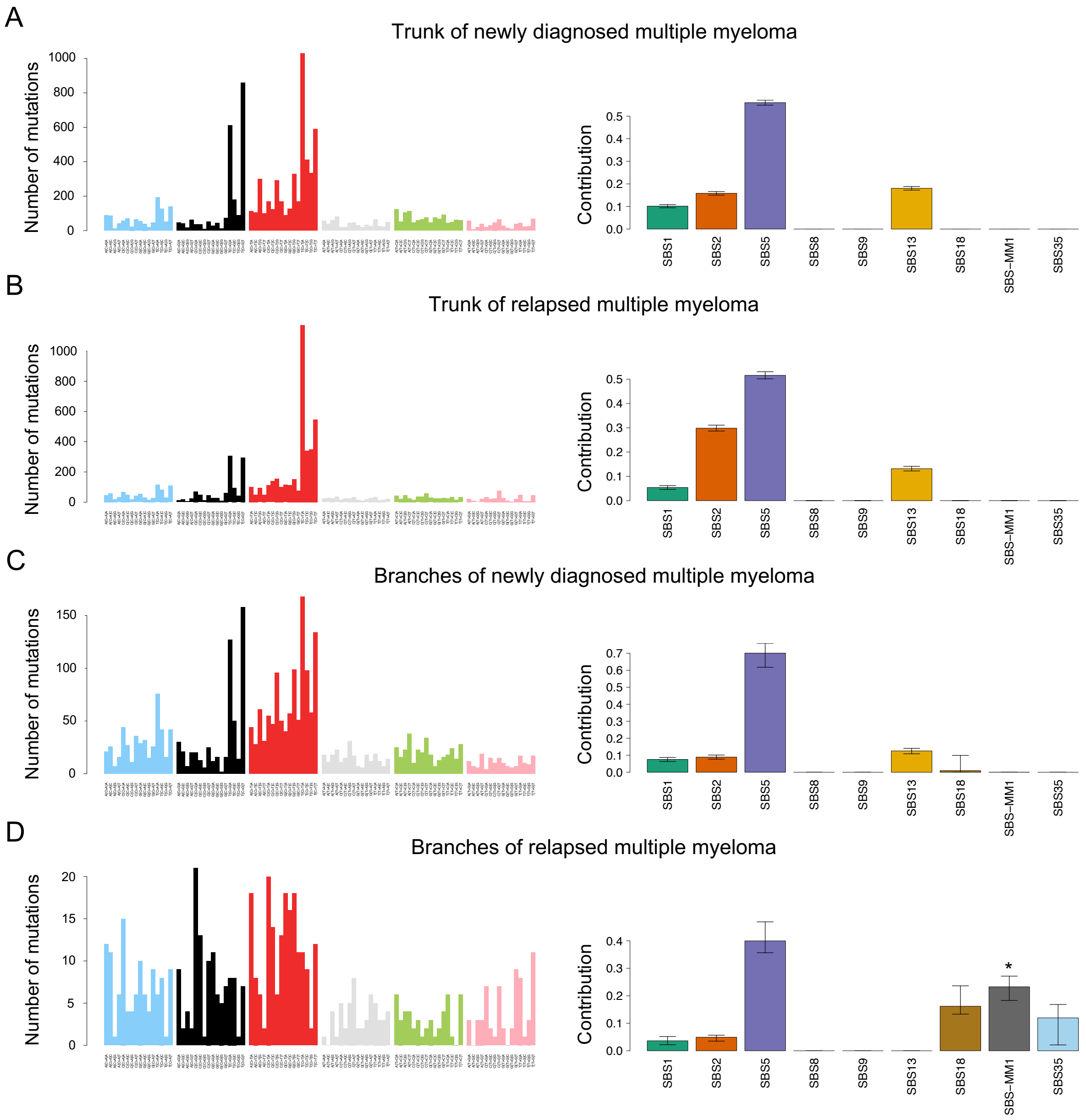

**Supplementary Figure 8.** Difference in SBS1 branches:trunk ratio between WXS at diagnosis, WGS and WXS relapsed multiple myeloma. p-value were estimated using Wilcoxon test.

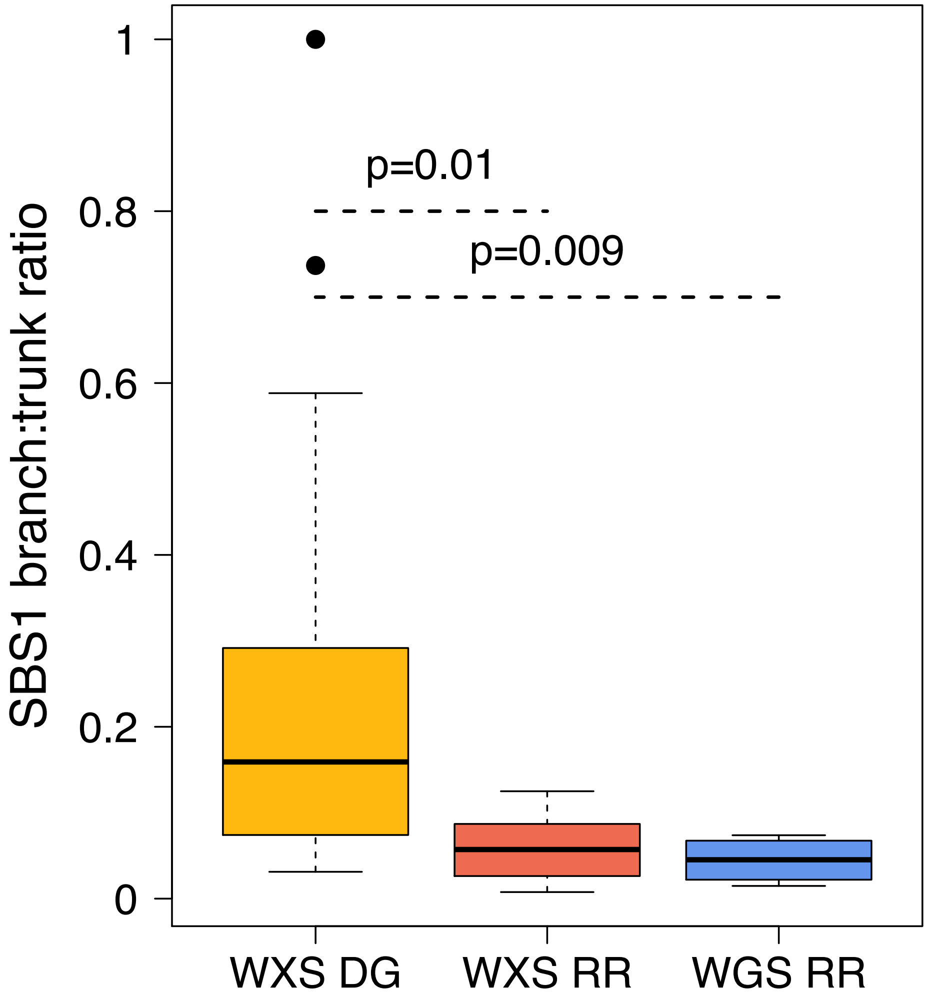

#### Supplementary Tables

**Supplementary Table 1.** WGS cohort demographic data.

| Patients | Gender | Isotype | Age |
|---|---|---|---|
| **I-H-106917** | Male | IgG Kappa | 53 |
| **I-H-130718** | Male | IgG Kappa | 60 |
| **I-H-130719** | Male | IgG Lambda | 54 |
| **I-H-130720** | Male | IgA Kappa | 64 |

**Supplementary Table 2.** Coverage, purity and ploidy of the WGS cohort.

| Sample | Purity | Ploidy | Median Coverage |
|---|---|---|---|
| **I-H-106917-T2-1-D1-2** | 0.6678 | 2.53 | 88.3745 |
| **I-H-106917-T2-2-D1-2** | 0.76726185 | 2.89423837 | 129.051 |
| **I-H-106917-T2-3-D1-2** | 0.78763 | 2.6048 | 101.449 |
| **I-H-106917-T2-4-D1-2** | 0.77841 | 2.4564 | 145.224 |
| **I-H-130718-T1-1-D1-2** | 0.94771604 | 2.15905083 | 89.9814 |
| **I-H-130718-T1-10-D1-2** | 0.9108 | 2.15 | 88.266 |
| **I-H-130718-T1-11-D1-1** | 0.74708 | 2.1824 | 121.605 |
| **I-H-130718-T1-12-D1-1** | 0.7 | 2.25 | 132.268 |
| **I-H-130718-T1-2-D1-2** | 0.92814 | 2.15 | 98.4156 |
| **I-H-130718-T1-4-D1-2** | 0.92 | 2.15 | 87.7814 |
| **I-H-130718-T1-6-D1-2** | 0.96419548 | 2.24056118 | 89.4443 |
| **I-H-130718-T1-9-D1-2** | 0.82908 | 2.2 | 89.8941 |
| **I-H-130719-T1-2-D1-2** | 0.8159 | 2.35 | 109.137 |
| **I-H-130719-T1-4-D1-2** | 0.2755 | 2.4 | 92.357 |
| **I-H-130719-T1-5-D1-2** | 0.69156 | 2.35 | 89.085 |
| **I-H-130719-T1-6-D1-2** | 0.68136 | 2.4 | 99.8962 |
| **I-H-130720-T1-2-D1-2** | 0.97924447 | 2.04353711 | 91.9502 |
| **I-H-130720-T1-3-D1-2** | 0.9598966 | 2.01730807 | 93.0744 |
| **I-H-130720-T1-4-D1-2** | 0.92907 | 2.05 | 88.1103 |
| **I-H-130720-T1-5-D1-2** | 0.97575174 | 2.05223504 | 88.2675 |
| **I-H-130720-T1-8-D1-2** | 0.95006146 | 2.02236114 | 88.3816 |
| **I-H-130720-T1-9-D1-1** | 0.26769043 | 2.8866776 | 107.828 |

**Supplementary Table 3.** Clinical and spatial characteristic of the whole exome cohort.

| Study ID | Sample | Stage | Platinum-based | Melphalan-based | Posterior iliac crest | Biopsy |
|---|---|---|---|---|---|---|
| **1** | IID_H107686 | Diagnosis | - | - | left | L4 |
| **2** | IID_H107681 | Diagnosis | - | - | left | Ilium |
| **3** | IID_H107677 | Diagnosis | - | - | right | Symphysis |
| **4** | IID_H107675 | Diagnosis | - | - | right | Ilium |
| **5** | IID_H107685 | Diagnosis | - | - | right | Ilium |
| **6** | IID_H107683 | Diagnosis | - | - | left | Sacrum |
| **7** | IID_H107679 | Diagnosis | - | - | right | T8, Ilium |
| **8** | IID_H107682 | Diagnosis | - | - | left | Rib, Pelvis, L1 |
| **9** | IID_H107678 | Diagnosis | - | - | left | Ilium |
| **10** | IID_H107684 | Diagnosis | - | - | right | L1 |
| **11** | IID_H107674 | Diagnosis | - | - | left | L4 |
| **12** | IID_H107676 | Diagnosis | - | - | left | T5, Sacrum, Ilium x 2 |
| **13** | IID_H107680 | Diagnosis | - | - | right | T8 |
| **14** | IID_H159193 | Diagnosis | - | - | right | Iliac |
| **15** | IID_H159194 | Diagnosis | - | - | right | Sacrum, Sacrum |
| **16** | IID_H159195 | Diagnosis | - | - | right | T5 |
| **17** | IID_H159196 | Diagnosis | - | - | left | Acetabulum |
| **18** | IID_H159197 | Diagnosis | - | - | left | Ilium |
| **19** | IID_H159198 | Diagnosis | - | - | right | Clavicle |
| **20** | IID_H159199 | Diagnosis | - | - | right | T12, T8, Acetabulum |
| **21** | IID_H159200 | Diagnosis | - | - | left | L3 |
| **22** | IID_H159201 | Diagnosis | - | - | left | L5 |
| **23** | IID_H159202 | Diagnosis | - | - | left | Sacrum |
| **24** | IID_H159203 | Diagnosis | - | - | right | T7 |
| **25** | IID_H159204 | Diagnosis | - | - | left | T8, L2, L3 |
| **26** | IID_H159205 | Diagnosis | - | - | left | Sacrum |
| **27** | IID_H159206 | Diagnosis | - | - | left | L1 |
| **28** | IID_H159207 | Diagnosis | - | - | left | Ischium, Ilium |
| **28** | IID_H159207 | Diagnosis | no | no | right | Pelvis |
| **29** | IID_H159208 | Diagnosis | - | - | left | Sacrum |
| **30** | IID_H159209 | Diagnosis | - | - | left | Anterior |
| **31** | IID_H159210 | Diagnosis | - | - | left | Ilium, T10, L1 |
| **48** | IID_H159227 | Diagnosis | no | no | left | Ischium, Ilium, T8 |
| **32** | IID_H159211 | Diagnosis | - | - | right | Sacrum |
| **33** | IID_H159212 | Diagnosis | - | - | right | Sacrum |
| **34** | IID_H159213 | Diagnosis | - | - | right | Pleural |
| **35** | IID_H159214 | Diagnosis | - | - | left | Posterior |
| **38** | IID_H159217 | Diagnosis | - | - | right | Sacrum |
| **39** | IID_H159218 | Diagnosis | - | - | right | Sacrum |
| **40** | IID_H159219 | Diagnosis | - | - | right | Ilium |
| **41** | IID_H159220 | Diagnosis | - | - | left | T8 |
| **42** | IID_H159221 | Diagnosis | - | - | left | Posterior |
| **45** | IID_H159224 | Relapse | yes | no | - | T3 x2 |
| **32** | IID_H159211 | Relapse | yes | yes | right | Sacrum |
| **43** | IID_H159222 | Relapse | yes | yes | right | Sacrum, T9 |
| **44** | IID_H159223 | Relapse | yes | yes | right | Sacrum |

| | | | | | | |
|---|---|---|---|---|---|---|
| **46** | IID_H159225 | Relapse | yes | yes | left | L4 |
| **47** | IID_H159226 | Relapse | yes | yes | left | Sacrum |
| **49** | IID_H159228 | Relapse | yes | yes | left | Right |
| **50** | IID_H159229 | Relapse | yes | yes | right | L2, Ilium |
| **51** | IID_H159192 | Relapse | yes | yes | right | T12 |

**Supplementary Table 4.** Coverage and purity characteristics of the whole exome cohort.

| Sample | Patients | Study ID | Purity | Ploidy | Median Coverage |
|---|---|---|---|---|---|
| **IID_H107674_T05_01_WE01** | IID_H107674 | 11 | 0.8002272 | 2.2694711 | 128 |
| **IID_H107674_T06_01_WE01** | IID_H107674 | 11 | 0.6091187 | 2.2481825 | 116 |
| **IID_H107675_T05_01_WE01** | IID_H107675 | 4 | 0.60931558 | 1.86780442 | 159 |
| **IID_H107675_T06_01_WE01** | IID_H107675 | 4 | 0.50668536 | 1.93331034 | 99 |
| **IID_H107676_T05_01_WE01** | IID_H107676 | 12 | 0.97001951 | 2.09682618 | 88 |
| **IID_H107676_T06_01_WE01** | IID_H107676 | 12 | 0.96428541 | 2.09482283 | 82 |
| **IID_H107676_T07_01_WE01** | IID_H107676 | 12 | 0.94218106 | 2.13657702 | 70 |
| **IID_H107676_T08_01_WE01** | IID_H107676 | 12 | 0.95948579 | 2.12620523 | 87 |
| **IID_H107676_T09_01_WE01** | IID_H107676 | 12 | 0.50818037 | 2.20878185 | 82 |
| **IID_H107677_T05_01_WE01** | IID_H107677 | 3 | 0.8665444 | 2.05411238 | 73 |
| **IID_H107677_T06_01_WE01** | IID_H107677 | 3 | 0.74822104 | 2.01652858 | 124 |
| **IID_H107678_T05_01_WE01** | IID_H107678 | 9 | 0.94894169 | 1.97271947 | 80 |
| **IID_H107678_T06_01_WE01** | IID_H107678 | 9 | 0.89273631 | 1.97978789 | 82 |
| **IID_H107679_T05_01_WE01** | IID_H107679 | 7 | 0.5838939 | 2.3652043 | 59 |
| **IID_H107679_T06_01_WE01** | IID_H107679 | 7 | 0.8354561 | 2.3706632 | 62 |
| **IID_H107679_T07_01_WE01** | IID_H107679 | 7 | 0.8047266 | 2.3729548 | 62 |
| **IID_H107680_T05_01_WE01** | IID_H107680 | 13 | 0.8438806 | 2.1559941 | 86 |
| **IID_H107680_T06_01_WE01** | IID_H107680 | 13 | 0.8333836 | 2.1744959 | 91 |
| **IID_H107681_T05_01_WE01** | IID_H107681 | 2 | 0.9630114 | 2.3173836 | 98 |
| **IID_H107681_T06_01_WE01** | IID_H107681 | 2 | 0.6284588 | 2.5577561 | 103 |
| **IID_H107682_T05_01_WE01** | IID_H107682 | 8 | 0.8939267 | 2.177801 | 82 |
| **IID_H107682_T06_01_WE01** | IID_H107682 | 8 | 0.921053 | 2.1734415 | 83 |
| **IID_H107682_T07_01_WE01** | IID_H107682 | 8 | 0.8197613 | 2.176688 | 125 |
| **IID_H107682_T08_01_WE01** | IID_H107682 | 8 | 0.9217819 | 2.2564858 | 127 |
| **IID_H107683_T04_01_WE01** | IID_H107683 | 6 | 0.7353756 | 2.5173613 | 110 |
| **IID_H107683_T05_01_WE01** | IID_H107683 | 6 | 0.7644068 | 2.5169572 | 136 |
| **IID_H107684_T05_01_WE01** | IID_H107684 | 10 | 0.73348935 | 1.9112504 | 90 |
| **IID_H107684_T06_01_WE01** | IID_H107684 | 10 | 0.94217719 | 1.99039462 | 65 |
| **IID_H107685_T05_01_WE01** | IID_H107685 | 5 | 0.8603543 | 2.6715024 | 111 |
| **IID_H107685_T06_01_WE01** | IID_H107685 | 5 | 0.560074 | 2.3485615 | 73 |
| **IID_H107686_T05_01_WE01** | IID_H107686 | 1 | 0.94766558 | 1.93627571 | 110 |
| **IID_H107686_T06_01_WE01** | IID_H107686 | 1 | 0.6659103 | 2.2234326 | 100 |
| **IID_H159192_T01_01_WE01** | IID_H159192 | 51 | 0.83804261 | 2.07896957 | 95 |
| **IID_H159192_T02_01_WE01** | IID_H159192 | 51 | 0.82906367 | 2.08565467 | 95 |
| **IID_H159193_T01_01_WE01** | IID_H159193 | 14 | 0.2853044 | 1.6461001 | 77 |
| **IID_H159193_T02_01_WE01** | IID_H159193 | 14 | 0.2793778 | 1.70384482 | 77 |
| **IID_H159194_T01_01_WE01** | IID_H159194 | 15 | 0.949149 | 2.1787565 | 71 |
| **IID_H159194_T02_01_WE01** | IID_H159194 | 15 | 0.97119946 | 2.13396304 | 91 |

| | | | | | |
|---|---|---|---|---|---|
| **IID_H159194_T03_01_WE01** | IID_H159194 | 15 | 0.90459305 | 2.14247739 | 79 |
| **IID_H159195_T01_01_WE01** | IID_H159195 | 16 | 0.95128944 | 2.01012549 | 66 |
| **IID_H159195_T02_01_WE01** | IID_H159195 | 16 | 0.93960744 | 1.99212335 | 85 |
| **IID_H159196_T01_01_WE01** | IID_H159196 | 17 | 0.8384426 | 2.3350139 | 105 |
| **IID_H159196_T02_01_WE01** | IID_H159196 | 17 | 0.4478289 | 2.29280087 | 84 |
| **IID_H159197_T01_01_WE01** | IID_H159197 | 18 | 0.9224657 | 2.5700855 | 98 |
| **IID_H159197_T02_01_WE01** | IID_H159197 | 18 | 0.8507106 | 2.539911 | 114 |
| **IID_H159198_T01_01_WE01** | IID_H159198 | 19 | 0.94679 | 2.3540743 | 90 |
| **IID_H159198_T02_01_WE01** | IID_H159198 | 19 | 0.8608714 | 2.3666977 | 90 |
| **IID_H159199_T01_01_WE01** | IID_H159199 | 20 | 0.7013097 | 2.400672 | 96 |
| **IID_H159199_T02_01_WE01** | IID_H159199 | 20 | 0.8488204 | 2.3950974 | 97 |
| **IID_H159199_T03_01_WE01** | IID_H159199 | 20 | 0.9053691 | 2.3947028 | 82 |
| **IID_H159199_T04_01_WE01** | IID_H159199 | 20 | 0.8603973 | 2.3947498 | 94 |
| **IID_H159200_T01_01_WE01** | IID_H159200 | 21 | 0.96572423 | 2.04134535 | 133 |
| **IID_H159200_T02_01_WE01** | IID_H159200 | 21 | 0.86389517 | 2.04768843 | 131 |
| **IID_H159201_T01_01_WE01** | IID_H159201 | 22 | 0.49243245 | 1.99047269 | 143 |
| **IID_H159201_T02_01_WE01** | IID_H159201 | 22 | 0.76782059 | 1.97094179 | 153 |
| **IID_H159202_T01_01_WE01** | IID_H159202 | 23 | 0.8718508 | 2.206232 | 89 |
| **IID_H159202_T02_01_WE01** | IID_H159202 | 23 | 0.7162207 | 2.2211288 | 123 |
| **IID_H159203_T01_01_WE01** | IID_H159203 | 24 | 0.7806844 | 2.4000219 | 91 |
| **IID_H159203_T02_01_WE01** | IID_H159203 | 24 | 0.8748083 | 2.4049286 | 95 |
| **IID_H159204_T01_01_WE01** | IID_H159204 | 25 | 0.662403 | 2.230165 | 105 |
| **IID_H159204_T02_01_WE01** | IID_H159204 | 25 | 0.68983017 | 2.19421531 | 133 |
| **IID_H159204_T03_01_WE01** | IID_H159204 | 25 | 0.7695526 | 2.196245 | 126 |
| **IID_H159204_T04_01_WE01** | IID_H159204 | 25 | 0.6537956 | 2.223384 | 117 |
| **IID_H159205_T01_01_WE01** | IID_H159205 | 26 | 0.1108276 | 2.6798439 | 75 |
| **IID_H159205_T02_01_WE01** | IID_H159205 | 26 | 0.95525302 | 2.06748533 | 78 |
| **IID_H159206_T01_01_WE01** | IID_H159206 | 27 | 0.6056648 | 2.3623408 | 65 |
| **IID_H159206_T02_01_WE01** | IID_H159206 | 27 | 0.7959656 | 2.3905351 | 77 |
| **IID_H159207_T01_01_WE01** | IID_H159207 | 28 | 0.7339352 | 4.2380534 | 114 |
| **IID_H159207_T01_01_WE01** | IID_H159207 | 28 | 0.7339352 | 4.2380534 | 114 |
| **IID_H159207_T02_01_WE01** | IID_H159207 | 28 | 0.8440228 | 4.1297769 | 119 |
| **IID_H159207_T02_01_WE01** | IID_H159207 | 28 | 0.8440228 | 4.1297769 | 119 |
| **IID_H159207_T03_01_WE01** | IID_H159207 | 28 | 0.7653815 | 2.4118943 | 141 |
| **IID_H159207_T03_01_WE01** | IID_H159207 | 28 | 0.7653815 | 2.4118943 | 141 |
| **IID_H159207_T04_01_WE01** | IID_H159207 | 28 | 0.5479146 | 4.457577 | 103 |
| **IID_H159207_T04_01_WE01** | IID_H159207 | 28 | 0.5479146 | 4.457577 | 103 |
| **IID_H159207_T05_01_WE01** | IID_H159207 | 28 | 0.4643885 | 2.5469944 | 118 |
| **IID_H159207_T05_01_WE01** | IID_H159207 | 28 | 0.4643885 | 2.5469944 | 118 |
| **IID_H159208_T01_01_WE01** | IID_H159208 | 29 | 0.7877964 | 2.00249734 | 80 |
| **IID_H159208_T02_01_WE01** | IID_H159208 | 29 | 0.79442943 | 1.9885857 | 84 |
| **IID_H159209_T01_01_WE01** | IID_H159209 | 30 | 0.9637269 | 2.1989709 | 114 |

| | | | | | |
|---|---|---|---|---|---|
| **IID_H159209_T02_01_WE01** | IID_H159209 | 30 | 0.7879392 | 2.1929875 | 126 |
| **IID_H159210_T01_01_WE01** | IID_H159210 | 31 | 0.92712086 | 2.00858273 | 83 |
| **IID_H159210_T02_01_WE01** | IID_H159210 | 31 | 0.93806964 | 1.97343159 | 107 |
| **IID_H159210_T03_01_WE01** | IID_H159210 | 31 | 0.91711508 | 1.99323632 | 98 |
| **IID_H159210_T04_01_WE01** | IID_H159210 | 31 | 0.86551271 | 1.96390226 | 120 |
| **IID_H159211_T01_01_WE01** | IID_H159211 | 32 | 0.8783528 | 2.03188819 | 137 |
| **IID_H159211_T01_01_WE01** | IID_H159211 | 32 | 0.8783528 | 2.03188819 | 137 |
| **IID_H159211_T02_01_WE01** | IID_H159211 | 32 | 0.63023737 | 2.04069739 | 131 |
| **IID_H159211_T02_01_WE01** | IID_H159211 | 32 | 0.63023737 | 2.04069739 | 131 |
| **IID_H159211_T03_01_WE01** | IID_H159211 | 32 | 0.56188275 | 2.01511523 | 176 |
| **IID_H159211_T03_01_WE01** | IID_H159211 | 32 | 0.56188275 | 2.01511523 | 176 |
| **IID_H159211_T04_01_WE01** | IID_H159211 | 32 | 0.81070812 | 2.01084752 | 104 |
| **IID_H159211_T04_01_WE01** | IID_H159211 | 32 | 0.81070812 | 2.01084752 | 104 |
| **IID_H159212_T01_01_WE01** | IID_H159212 | 33 | 0.8652078 | 2.2682235 | 114 |
| **IID_H159212_T02_01_WE01** | IID_H159212 | 33 | 0.5051255 | 2.27083046 | 115 |
| **IID_H159213_T01_01_WE01** | IID_H159213 | 34 | 0.96980921 | 1.96990208 | 102 |
| **IID_H159213_T02_01_WE01** | IID_H159213 | 34 | 0.75708204 | 1.97584576 | 125 |
| **IID_H159214_T01_01_WE01** | IID_H159214 | 35 | 0.7750537 | 2.4587212 | 75 |
| **IID_H159214_T02_01_WE01** | IID_H159214 | 35 | 0.7905361 | 2.461937 | 73 |
| **IID_H159214_T03_01_WE01** | IID_H159214 | 35 | 0.7797573 | 2.4679043 | 70 |
| **IID_H159215_T01_01_WE01** | IID_H159215 | 36 | 0.92918503 | 2.01415484 | 98 |
| **IID_H159215_T02_01_WE01** | IID_H159215 | 36 | 0.69855941 | 2.01105364 | 99 |
| **IID_H159216_T01_01_WE01** | IID_H159216 | 37 | 0.8330706 | 2.1822886 | 129 |
| **IID_H159216_T02_01_WE01** | IID_H159216 | 37 | 0.82611333 | 2.15428486 | 135 |
| **IID_H159217_T01_01_WE01** | IID_H159217 | 38 | 0.716789 | 1.9613428 | 133 |
| **IID_H159217_T02_01_WE01** | IID_H159217 | 38 | 0.6839017 | 2.0669228 | 107 |
| **IID_H159218_T01_01_WE01** | IID_H159218 | 39 | 0.94526328 | 1.93725482 | 88 |
| **IID_H159218_T02_01_WE01** | IID_H159218 | 39 | 0.93183277 | 1.93351012 | 84 |
| **IID_H159219_T01_01_WE01** | IID_H159219 | 40 | 0.89509 | 1.8119822 | 77 |
| **IID_H159219_T02_01_WE01** | IID_H159219 | 40 | 0.572952 | 1.82642918 | 113 |
| **IID_H159220_T01_01_WE01** | IID_H159220 | 41 | 0.8937164 | 2.2900012 | 99 |
| **IID_H159220_T02_01_WE01** | IID_H159220 | 41 | 0.8173879 | 2.2983591 | 95 |
| **IID_H159221_T01_01_WE01** | IID_H159221 | 42 | 0.8370796 | 2.6851918 | 117 |
| **IID_H159221_T02_01_WE01** | IID_H159221 | 42 | 0.9003912 | 2.7185433 | 110 |
| **IID_H159222_T01_01_WE01** | IID_H159222 | 43 | 0.743228 | 3.047736 | 77 |
| **IID_H159222_T02_01_WE01** | IID_H159222 | 43 | 0.7252859 | 3.1409718 | 79 |
| **IID_H159222_T03_01_WE01** | IID_H159222 | 43 | 0.79253508 | 1.86817258 | 84 |
| **IID_H159223_T01_01_WE01** | IID_H159223 | 44 | 0.96868429 | 2.11916229 | 95 |
| **IID_H159223_T02_01_WE01** | IID_H159223 | 44 | 0.47157677 | 2.22985945 | 77 |
| **IID_H159224_T01_01_WE01** | IID_H159224 | 45 | 0.8738939 | 2.609495 | 66 |
| **IID_H159224_T02_01_WE01** | IID_H159224 | 45 | 0.7960685 | 4.6081039 | 95 |
| **IID_H159225_T01_01_WE01** | IID_H159225 | 46 | 0.5272777 | 1.9398966 | 102 |

| | | | | | |
|---|---|---|---|---|---|
| **IID_H159225_T02_01_WE01** | IID_H159225 | 46 | 0.47170094 | 1.93603211 | 120 |
| **IID_H159226_T01_01_WE01** | IID_H159226 | 47 | 0.92537319 | 1.99920882 | 96 |
| **IID_H159226_T02_01_WE01** | IID_H159226 | 47 | 0.53255266 | 1.91373547 | 123 |
| **IID_H159227_T01_01_WE01** | IID_H159227 | 48 | 0.9107675 | 2.3705624 | 117 |
| **IID_H159227_T02_01_WE01** | IID_H159227 | 48 | 0.9751333 | 2.3660556 | 115 |
| **IID_H159227_T03_01_WE01** | IID_H159227 | 48 | 0.9383707 | 2.3171541 | 129 |
| **IID_H159227_T04_01_WE01** | IID_H159227 | 48 | 0.7306523 | 2.3878758 | 115 |
| **IID_H159228_T01_01_WE01** | IID_H159228 | 49 | 0.9660542 | 2.3084885 | 103 |
| **IID_H159228_T02_01_WE01** | IID_H159228 | 49 | 0.8801174 | 2.2496426 | 96 |
| **IID_H159229_T01_01_WE01** | IID_H159229 | 50 | 0.8335068 | 2.2233683 | 67 |
| **IID_H159229_T02_01_WE01** | IID_H159229 | 50 | 0.9234834 | 2.2469325 | 66 |
| **IID_H159229_T03_01_WE01** | IID_H159229 | 50 | 0.8911536 | 2.2505918 | 67 |

**Supplementary Table 5** *SigProfiler* assignment of mutational signatures.

| De novo extracted | Global NMF Signatures |
|---|---|
| **Signature 96-A** | Signature SBS9 (94.66%) |
| **Signature 96-B** | Signature SBS2 (64.42%) & Signature SBS13 (35.58%) |
| **Signature 96-C** | Signature SBS5 (57.54%) & Signature SBS35 (42.46%) |
| **Signature 96-D** | Signature SBS1 (17.80%) & Signature SBS5 (75.48%) |
| **Signature 96-E** | Signature SBS5 (63.62%) & Signature SBS9 (25.56%) |
| **Signature 96-F** | Signature SBS-MM1 |
| **Signature 96-G** | Signature SBS5 (27.52%) & Signature SBS8 (72.48%) |
| **Signature 96-H** | Signature SBS5 (47.20%) & Signature SBS18 (40.36%) |
